# A nanobody toolbox to investigate localisation and dynamics of *Drosophila* titins

**DOI:** 10.1101/2022.04.13.488177

**Authors:** Vincent Loreau, Renate Rees, Eunice HoYee Chan, Waltraud Taxer, Kathrin Gregor, Bianka Mußil, Christophe Pitaval, Nuno Miguel Luis, Pierre Mangeol, Frank Schnorrer, Dirk Görlich

**Affiliations:** Aix Marseille University, CNRS, IBDM, Turing Centre for Living Systems, 13288 Marseille, France; Max Planck Institute for Multidisciplinary Sciences, Göttingen, Germany

**Keywords:** nanobodies, titin, muscle, sarcomere, myofibril, *Drosophila*, FRAP

## Abstract

Measuring the positions and dynamics of proteins in intact tissues or whole animals is key to understand protein function. However, to date this is still a challenging task, as accessibility of large antibodies to dense tissues is often limited and fluorescent proteins inserted close to a domain of interest may affect function of the tagged protein. These complications are particularly present in the muscle sarcomere, arguably one of the most protein dense structures in nature, which makes studying morphogenesis at molecular resolution challenging. Here, we have employed an efficient pipeline to generate a nanobody toolbox specifically recognising various domains of two large *Drosophila* titin homologs, Sallimus and Projectin. We demonstrate the superior labelling qualities of our nanobodies compared to conventional antibodies in intact muscle tissue. Applying our nanobody toolbox to larval muscles revealed a gigantic Sallimus isoform stretched more than 2 µm to bridge the sarcomeric I-band. Furthermore, N- and C-terminal nanobodies against Projectin identified an unexpected polar orientation of Projectin covering the myosin filaments in larval muscles. Finally, expression of a Sallimus nanobody in living larval muscles confirmed the high affinity binding of nanobodies to target epitopes in living tissue and hence demonstrated their power to reveal the *in vivo* dynamics of sarcomeric protein domains. Together, our toolbox substantiates the multiple advantages of nanobodies to study sarcomere biology. It may inspire the generation of similar toolboxes for other large protein complexes in *Drosophila* or mammals.

## Introduction

Muscles use their sarcomeres to generate forces that power animal movements. The sarcomere morphologies are remarkably conserved from fruit flies to humans: each sarcomere is bordered by two Z-discs that cross-link the plus-ends of parallel actin filaments, while their minus ends face towards the centrally located bipolar myosin filaments (Lange et al., 2006; Lemke and Schnorrer, 2017). Both filaments are stably linked by the gigantic titin spring protein, which in mammals binds with its N-terminus to alpha-actinin at the Z-disc and is anchored with its C-terminus at the M-band in the middle of the sarcomere. Such, titin determines the length of the mammalian sarcomere (Linke, 2018; Luis and Schnorrer, 2021; Tskhovrebova and Trinick, 2003).

As muscle and in particular sarcomere architecture is well-conserved, *Drosophila* is a fantastic model to study how a sarcomere is built during development (Katzemich et al., 2013; 2015; Orfanos et al., 2015; Weitkunat et al., 2017; 2014). Generation of monoclonal antibodies against *Drosophila* sarcomere proteins have been insightful to locate key proteins within the mature sarcomere (Burkart et al., 2007; Ferguson et al., 1994; Katzemich et al., 2012; Lakey et al., 1990; Qiu et al., 2005; Szikora et al., 2020). However, a systematic toolbox of antibodies recognising defined domains of the often large sarcomeric proteins, in particular against defined domains of the two large *Drosophila* titin homologs Sallimus (Sls) and Projectin (gene called *bent, bt*) is still missing. These tools would be important to understand how the sarcomeric machine assembles during muscle morphogenesis.

Recent gene tagging approaches have generated a substantial amount of *Drosophila* transgenic lines, each expressing one sarcomeric protein fused to GFP at its C-terminus (Sarov et al., 2016) or at a random internal position (Buszczak et al., 2007; Kanca et al., 2017; Kelso et al., 2004; Nagarkar-Jaiswal et al., 2015). Nevertheless, a number of these tagged lines label only a subset of protein isoforms or result in homozygous loss of function phenotypes as the GFP tagged protein cannot fully recapitulate the function of the endogenous protein in the dense sarcomeric environment (Orfanos and Sparrow, 2013; Orfanos et al., 2015; Sarov et al., 2016). Hence, classical GFP tagging does not always provide an optimal solution to study the native dynamics of a sarcomeric protein.

These limitations motivated us to develop a complementary toolbox to antibodies and GFP-tagged lines for sarcomeric proteins that will be particularly well suited for the dense environment of mature sarcomeres in intact muscles (O’Donnell et al., 1989). We chose the recent cameloid nanobody technology, because of the small size of nanobodies (<4 nm, 12- 15 kDa), their single chain protein nature and their high-affinity against target domains (Muyldermans, 2013; Pleiner et al., 2018; 2015). As nanobodies can be used on fixed samples or fused to a fluorescent protein and expressed in living tissues, nanobodies are ideal tools to quantify the position and dynamics of sarcomeric proteins in their dense environment.

Thus far, the application of nanobodies to the *Drosophila* model was largely restricted to commercially available GFP and mCherry nanobodies that allowed to locate, trap or degrade GFP- or mCherry-tagged proteins in *Drosophila* tissue (Ákos et al., 2021; Caussinus et al., 2011; Harmansa and Affolter, 2018; Harmansa et al., 2017; 2015). Recently, nanobodies located *Drosophila* proteins tagged with short artificial nanotags (Xu et al., 2022). However, prior to our studies nanobodies binding to endogenous *Drosophila* protein domains had to our knowledge not yet been made.

Here we generated a nanobody toolbox against seven different epitopes of the two *Drosophila* titin homologs Sallimus (Sls) and Projectin (Proj). After recombinant expression and labelling, we verified their specificity as well as their superior penetration and labelling efficiencies compared to antibodies. Applying our nanobodies to *Drosophila* muscle tissues confirmed the expression of different Sls isoforms in different muscle types and identified a gigantic more than 2 µm long Sls protein in larval muscles. It further revealed that Projectin is bound to the myosin filament in a strictly polar fashion, resembling the mammalian titin homolog. Finally, by generating transgenic animals expressing nanobody-Neongreen fusions, we verified that nanobodies are suitable tools to monitor the dynamics of endogenous sarcomeric proteins in intact living animals.

## Results

### *Drosophila* titin nanobody design

Mammalian sarcomere length is determined by a long titin protein isoform that spans linearly from the Z-disc to the M-band and thus adopts a length of about 1.5 µm in relaxed human muscle (Brynnel et al., 2018; Fürst et al., 1988; Linke, 2018). In order to investigate to what extend the localisation of this critical sarcomere component is conserved across evolution of bilaterian muscle, we aimed to re-investigate the localisation of the two *Drosophila* titin homologs Sallimus and Projectin by generating specific nanobodies. By carefully mining the Flybase expression database (http://flybase.org/reports/FBgn0086906; http://flybase.org/reports/FBgn0005666) we have annotated the likely longest Sallimus (Sls) and Projectin (Proj, gene called *bent, bt*) isoforms expressed in larval body wall muscles (Figure 1A, B). The longest Sls isoform contains 48 immunoglobulin (Ig)-domains of the total 52 Ig domains coded in the Sls gene (4 are selectively present in a short larval isoform). Additionally, Sls does contain long stretches of flexible regions rich in amino acids proline, valine, glutamic acid and lysine (PEVK) that form an elastic spring and five C-terminal fibronectin (Fn)-domains (Figure 1A) (Burkart et al., 2007). This domain organisation largely resembles the I-band part of mammalian titin (Tskhovrebova and Trinick, 2003).

**Figure 1.**
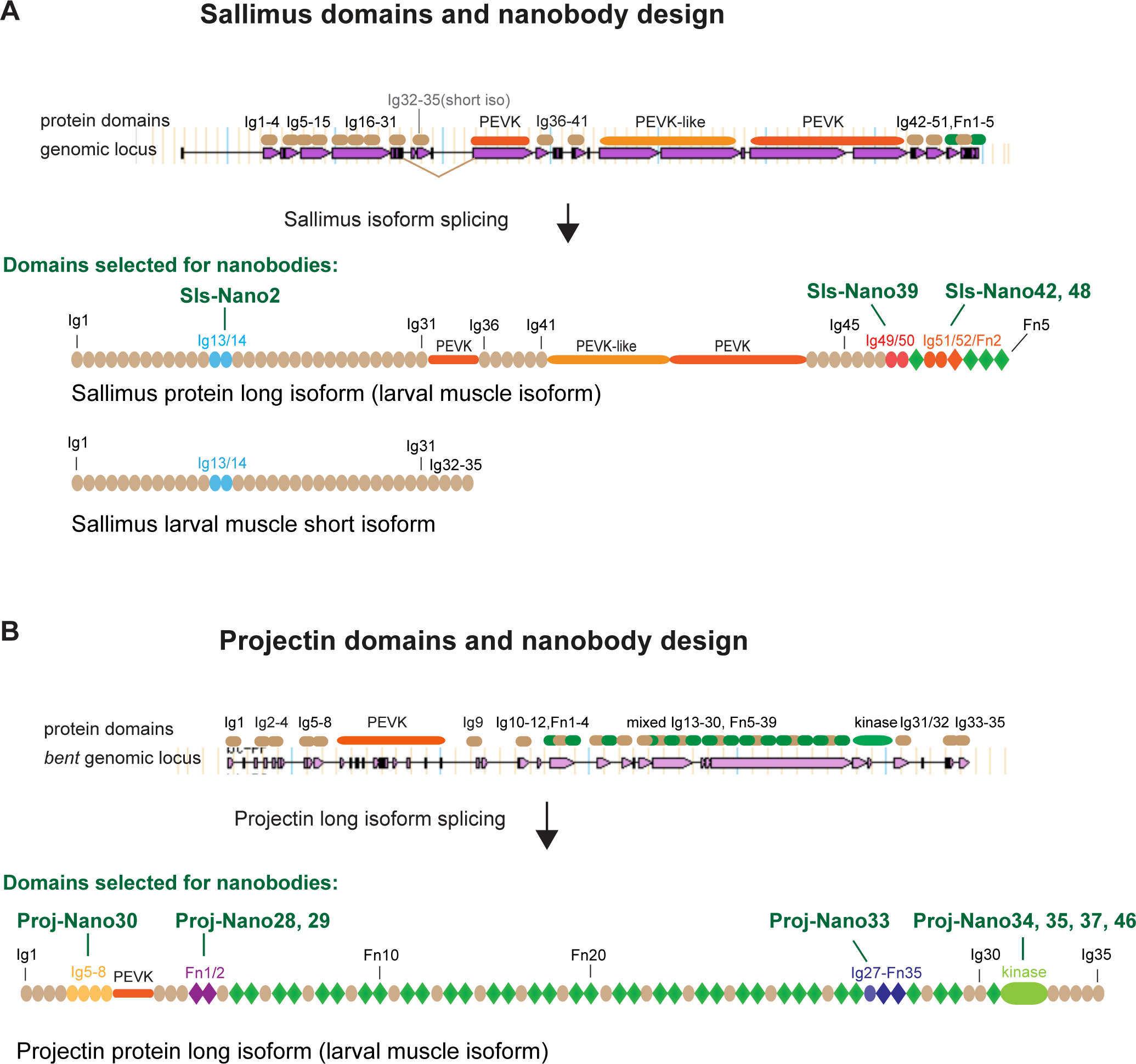
*Drosophila* titin nanobody design. (A, B) Schematic of Sallimus (A) and Projectin (B) gene and protein domain organisation. Top: genome loci taken from Flybase with exons represented by magenta arrows and introns by lines. The coded protein domains are overlayed with Ig-domains in brown, PEVK in orange, Fn- and kinase domains in green. Bottom: predicted domain organisation of Sls long and short (internal STOP) and Proj long larval muscle protein isoforms. Domains selected for nanobody production are highlighted by special colours and the names of the respective nanobodies are indicated above the protein.

Conversely, the long Projectin isoform contains 35 Ig- and 39 Fn-domains that are mainly organised in Ig-Fn super-repeats with a consensus myosin light chain kinase domain close to its C-terminus (Figure 1B) (Ayme-Southgate et al., 2008). This domain organisation largely resembles the A-band part of mammalian titin (Tskhovrebova and Trinick, 2003).

To generate nanobodies against Sls and Proj domains, we selected a subset of small domains that according to published transcriptomics data (Spletter et al., 2015; 2018) should also be expressed in other muscle types, such as flight or leg muscles. We chose domains close to the N- and C-termini of both proteins to assess their possible extended configuration in sarcomeres. We successfully expressed Sls-Ig13/14, Sls-Ig49/50 and SlsIg51/52 recombinantly and generated the respective nanobodies Sls-Nano2, Sls-Nano39, Sls-Nano42 and Sls-Nano48 against these domains (Figure 1A). For Projectin we selected Ig5-8, Fn1/2, Ig27-Fn35 as well as the kinase domain to generate Projectin nanobodies Proj-Nano30, Proj- Nano28 and 29, Proj-Nano33, and Proj-Nano34, 35, 37 and 46 recognising these domains, respectively (Figure 1B).

### *Drosophila* titin nanobody production

To effectively produce a comprehensive set of nanobodies against the above selected Sls and Proj domains, we used two sources of immunogens (Figure 2 for workflow). First, we hand-dissected the indirect flight muscles from 1000 wild-type adult flies and isolated their myofibrils, which express large amounts of Sls and Proj (Spletter et al., 2018). These were used for two immunisations of the alpaca. Second, we recombinantly expressed selected Sls and Proj domains as His_14_-SUMO or His_14_-NEDD8-tagged proteins in *E.coli* and purified them by binding to a Ni(II) chelate matrix, followed by extensive washing and elution with a tag-cleaving protease (Frey and Görlich, 2014a). These recombinant antigens (100 µg each) were used for three consecutive immunisations. Four days after the last immunisation, a blood sample was taken, lymphocytes were recovered, total RNA was isolated and reverse transcribed into cDNA. Finally, a phage display library with a complexity of more than 10^8^ independent clones was constructed. This followed a previously described workflow (Pleiner et al., 2015; 2018).

**Figure 2.**
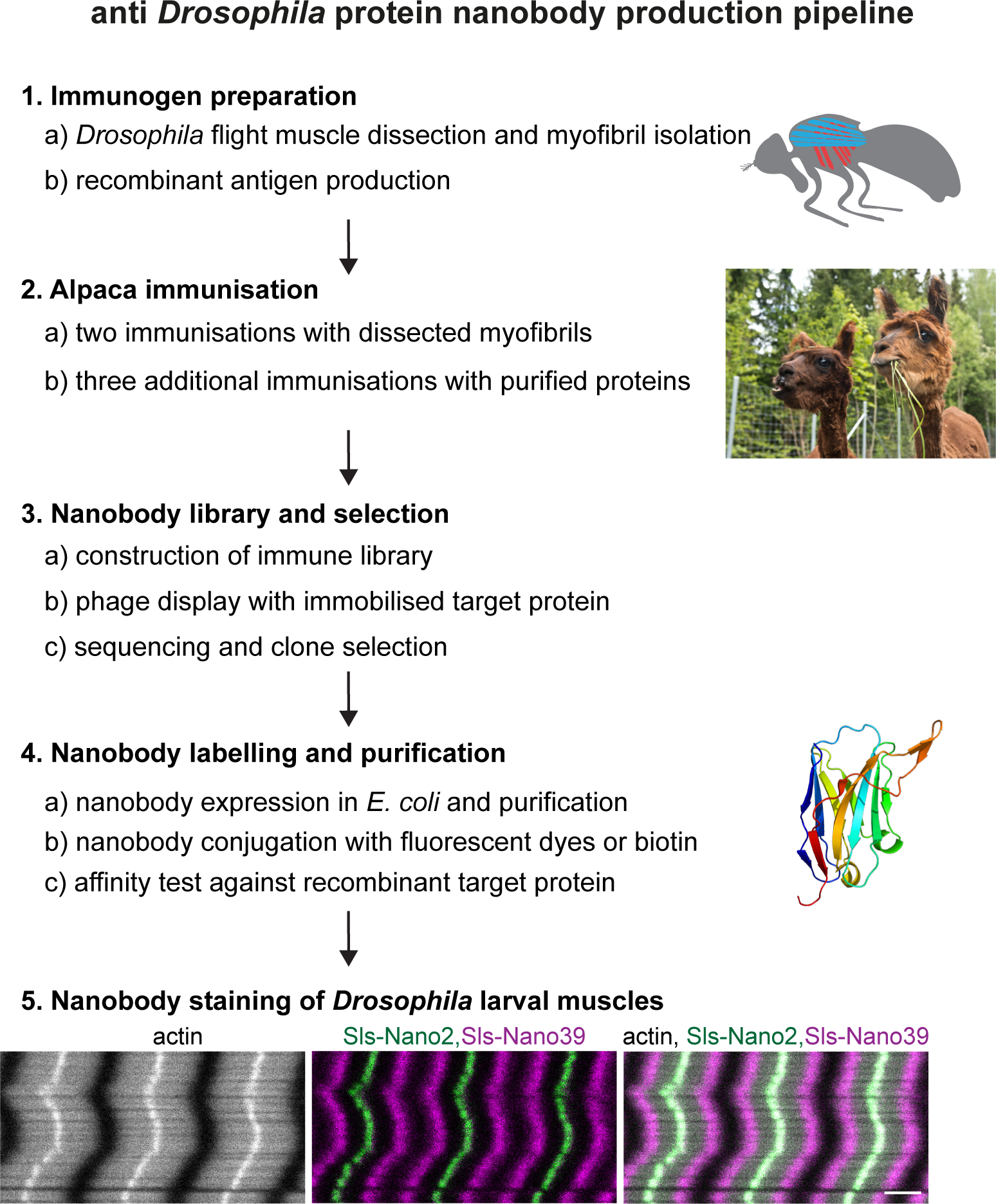
Nanobody production pipeline. Overview of our optimised nanobody production pipeline against *Drosophila* sarcomeric protein domains. See text for details. Scale bar is 3 µm.

To isolate high-affinity nanobodies, we employed three rounds of phage display, using low concentrations (1nM) of baits. Coding sequences of selected nanobodies were sequenced in a 96-well format and classified according to their sequence similarity. Selected clones were then expressed as His_14_-SUMO-tagged fusions in *E.coli* and purified by the affinity-capture- protease elution strategy, with typical yields of 20-50 mg nanobody per litre TB culture.

### Nanobody labelling and affinities

For application in fluorescence microscopy, we labelled the nanobodies directly through one or two ectopic cysteines (N- and C-terminal) with appropriate maleimides (Pleiner et al., 2015; 2018). The labelling was performed “on-column”, i.e., after binding the His_14_-SUMO-tagged nanobodies to Ni(II) chelate beads. Washing of the beads allowed for convenient removal of free fluorophore before the tag-free labelled nanobodies were eluted with the tag-cleaving protease. The labelling according to this workflow was nearly quantitative, as indicated by the observed size shifts on SDS-PAGE (Figure 3A) and by a ratiometric measurement of the optical densities at the absorption maxima of protein and the fluorophore.

**Figure 3.**
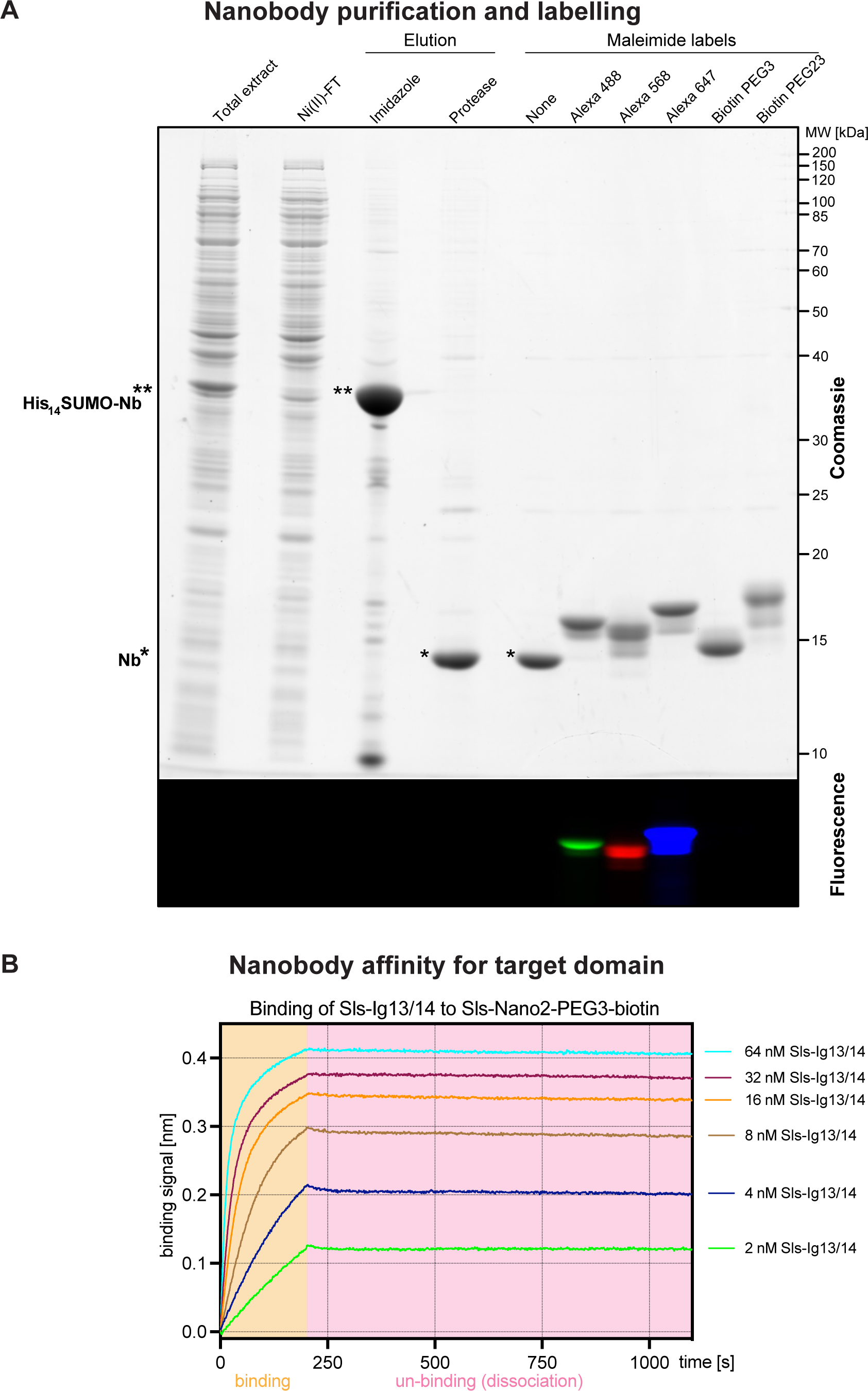
Nanobody labelling and affinity test. (**A**) SDS-PAGE gel documenting the expression, purification, tag cleavage and labelling of a nanobody, here Sls-Nano2. Top part stained with Coomassie blue, lower part shows fluorescence of the same gel. Note the efficiency of the labelling (essentially quantitative) by the size shift of the bands. (**B**) Nanobody affinity assay. Sls-Nano2-biotin was immobilised to high precision streptavidin Octet sensors to a binding signal of 0.4 nm. After washing, the target domain Sls-Ig13/14 was allowed to bind at indicated concentration for 200 seconds (beige box), followed by a 900 seconds dissociation step. A global fit of the curves indicates a 10 pM K_D_.

To measure the binding affinity (K_D_) of a nanobody to its target, we chose Sls-Nano2, labelled it with biotin, and immobilised it on high-precision streptavidin Octet sensors for biolayer interferometry (Figure 3B). On- and off-rates of the cognate Sls Ig13/14 domains were then measured by allowing a concentration series to bind and subsequently to dissociate from the nanobody (Figure 3B). The data indicate a nearly irreversible binding with an on-rate of ∼10^6^·M^-1^·sec^-1^, an off-rate in the order of 10^-5^·M^-1^·sec^-1^ and K_D_ in the 10 pM range. Note that such high affinity is already at the limit of what can be reliably measured with this technology. Such a high affinity can be explained by *Drosophila* proteins being highly immunogenic in mammals, by the very large immune repertoire of alpacas, and by our very stringent selection from a very large immune library.

### Sallimus and Projectin nanobody specificity

To assess the efficiency and specificity of our nanobodies in muscle tissue we first assayed how well they label late stage *Drosophila* embryonic muscles. We fixed wild type stage 17 embryos and incubated them with fluorescently labelled Sls or Proj nanobodies and performed confocal microscopy. We found that most of our nanobodies, which had passed our selection criteria (see Methods), efficiently stained embryonic muscles revealing the expected striated pattern of Sls and Proj in stage 17 embryos (Figure 4A, B and Figure 4 supplements 1 and 2). Thus, in total we generated 12 different Sls and Proj nanobodies against 3 different Sls and 4 different Proj epitopes.

**Figure 4.**
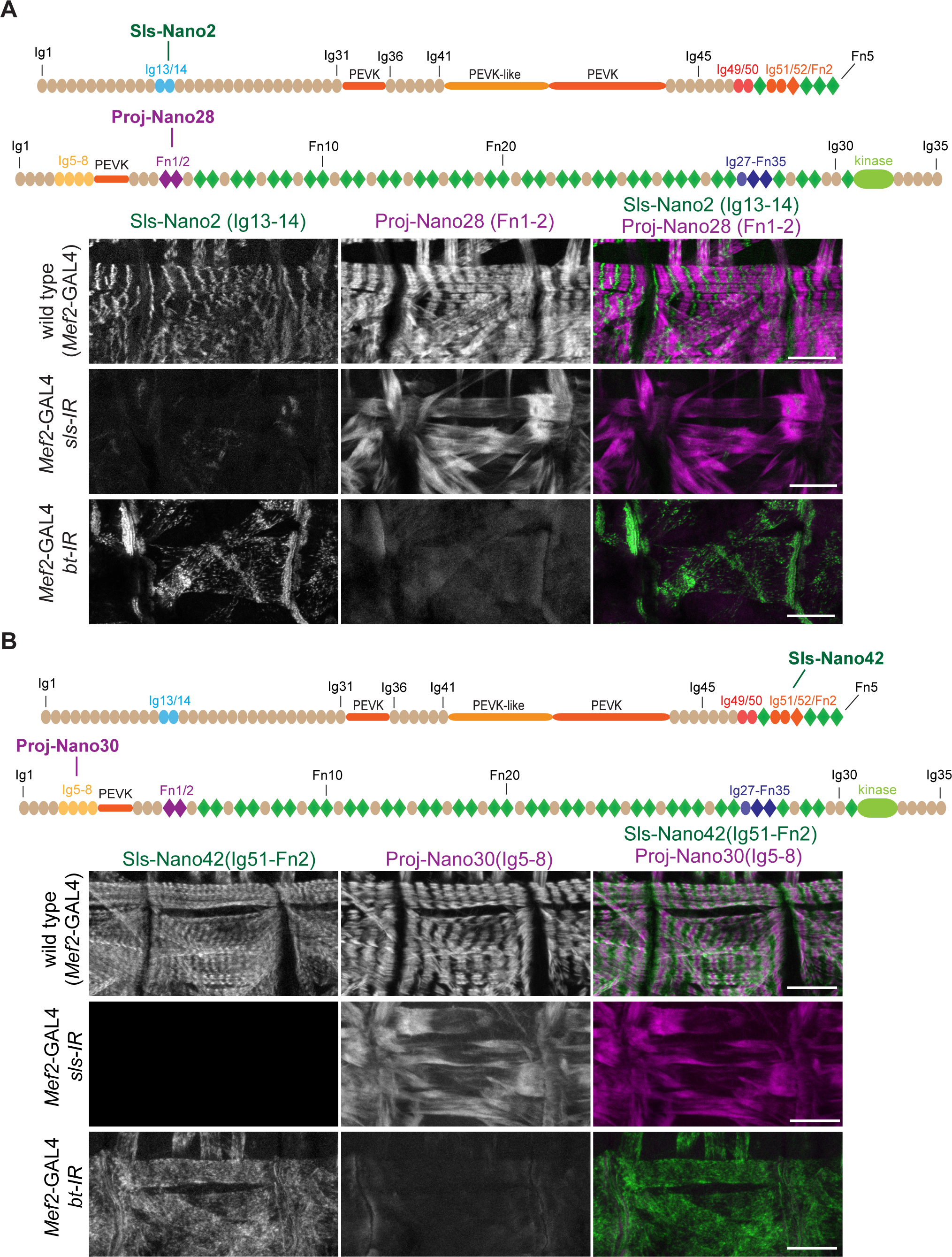
Sallimus and Projectin nanobody specificity. (**A, B**) top: schematic representation of Sallimus or Projectin domains with nanobodies used for stainings. Bottom: stage 17 embryos of wild type (*Mef2*-GAL4) and muscle specific *sls* or *bt* knock-down (*Mef2*-GAL4, UAS-*sls-IR* and *Mef2*-GAL4, UAS-*bt-IR*, respectively) stained with Sls (green) and Proj (magenta) nanobodies Sls-Nano2 and Proj-Nano28 (A) or Sls- Nano42 and Proj-Nano30 (B). Note the striated pattern of Sls and Proj in wild type, which is lost upon knock-down of one component. Scale bars are 20 µm.

To test for the specificity of the nanobodies we generated embryos in which we either depleted the Sls or Proj protein by muscle-specific RNAi using *Mef2*-GAL4 (Schnorrer et al., 2010), followed by a double staining with anti-Sls and anti-Proj nanobodies. Importantly, we found that in all cases the staining of Sls or Proj was severely reduced or lost entirely after knock-down of the respective protein, demonstrating the specificity of our nanobodies (Figure 4 and Figure 4 supplements 1 and 2). As expected, in each case we found that the striated pattern of the other protein was lost, demonstrating that both, Sls and Proj are required to generate striated sarcomeres in stage 17 embryos. We conclude that our nanobodies specifically recognise the various Sls and Proj domains against which they were raised and hence should be valuable tools to study the roles of the *Drosophila* titin homologs in sarcomere biology.

### Nanobodies display superior labelling and penetration efficiencies

Nanobodies are attractive labels because of their small size compared to conventional antibodies. We wanted to investigate if this proposed theoretical advantage matters in practice when staining *Drosophila* muscle samples. We compared the labelling of our nanobodies to conventional antibodies visualised with standard secondary antibodies in embryos at stage 16 and stage 17. The rational being that at stage 17, the future larval cuticle is already deposited around the embryos (Moussian, 2010), hindering the penetration of large labels into the embryos and thus rendering the staining ineffective when compared with stage 16. We used the nanobodies against Sls and Proj together with either Sls, myosin heavy chain (Mhc) or Proj antibodies (Figure 5A-C). In stage 16 embryos we found the expected co-localisation of Sls-Nano2, recognising Sls-Ig13/14, with an anti-Sls antibody, recognising Sls-Ig16 (called anti-Kettin) (Kulke et al., 2001) (Figure 5A), as well as the co-localisation of Proj-Nano30 recognising Proj-Ig5-8 with a Proj antibody (Figure 5C). Both Sls and Proj proteins are not yet displaying a striated pattern as sarcomeres have not yet assembled at stage 16.

**Figure 5.**
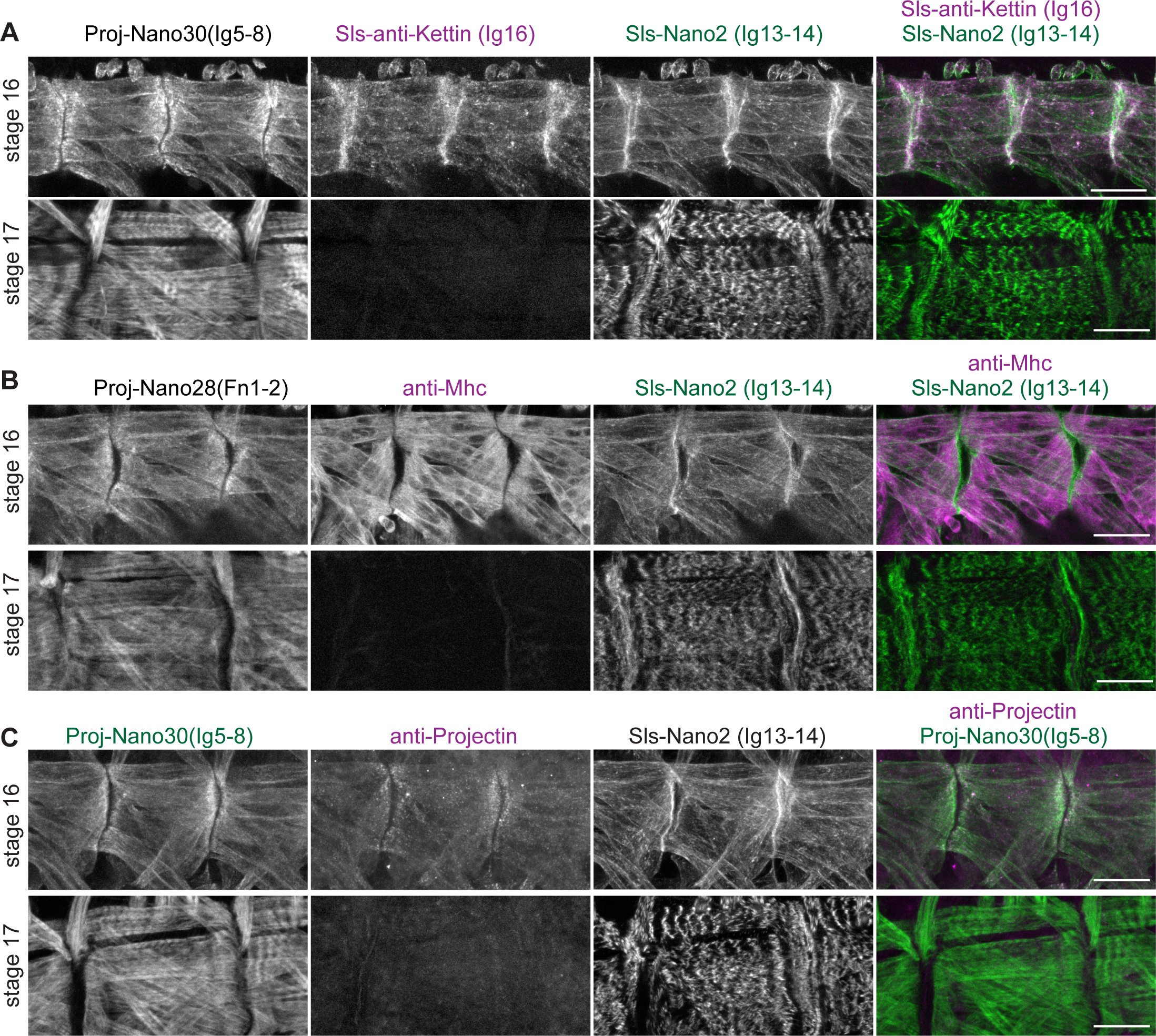
Nanobodies penetrate embryos easier than antibodies. (**A-C**) Stage 16 and 17 wild type embryos (*Mef2*-GAL4) stained with Sls and Proj nanobodies (Sls-Nano2, Proj-Nano28, Proj-Nano30) in green, in combination with antibodies against Sls (anti-Kettin) (A), Mhc (B) or Projectin (C) in magenta. Scale bar 20 µm. Note that Sls, Mhc and Proj antibodies cannot penetrate into stage 17 embryos efficiently, whereas the respective nanobodies stain muscles well.

Strikingly, while our nanobodies stained the body muscles of stage 17 embryos very well, displaying the striated pattern of the first formed sarcomeres, neither Kettin, Mhc nor Projectin antibodies produced a good staining pattern (Figure 5A-C). This demonstrates the superior penetration and labelling efficiency of small nanobodies compared to antibodies. We found similar differences when testing penetration efficiencies in large flight muscles in the accompanying manuscript (Schueder et al., 2022). Together, we conclude that the here generated Sls or Proj nanobody toolbox specifically recognises Sls or Proj domains, which allows efficient labelling of sarcomeres in whole mount late-stage embryos, i.e. at a stage that is impossible to investigate with antibodies following standard protocols.

### Sallimus and Projectin localisation in mature muscles

Our Sls and Proj toolbox should be highly valuable to investigate muscle development at various stages of the *Drosophila* life cycle as we had chosen Sls and Proj domains that are predicted to be expressed in most muscle types at most stages of development (see Figure 1). To test this prediction, we next investigated adult *Drosophila* flight muscles, which show a specialised fibrillar morphology of their myofibrils and sarcomeres that is caused by the expression of a specific combination of sarcomeric protein isoforms (Schönbauer et al., 2011; Spletter et al., 2015). Co-staining flight muscles with the Sls-Nano2 recognising Sls-Ig13/14 close to the N-term of Sls and Sls-Nano42 recognising Sls-Ig51-Fn2 close to the C-term of Sls revealed single and overlapping bands present at the sarcomeric Z-disc (Figure 6A). This pattern is expected since flight muscles contain a very short about 100 nm wide I-band (Burkart et al., 2007). The Sls-Nano42 band is shorter, with a smaller cross-sectional radius compared to Sls-Nano2 suggesting that Sls-Ig51-Fn2 is not present in all the Sls isoforms expressed during the final stages of myofibril maturation that complete radial myofibril growth (González-Morales et al., 2019; Spletter et al., 2018). Such a central localisation of the long Sls isoforms in flight muscle sarcomeres had also been reported previously with the anti-Sls antibody B2 that likely recognises Sls-Ig36-41 domains (Burkart et al., 2007), thus further confirming the specificity of our domain-specific Sls nanobodies.

**Figure 6.**
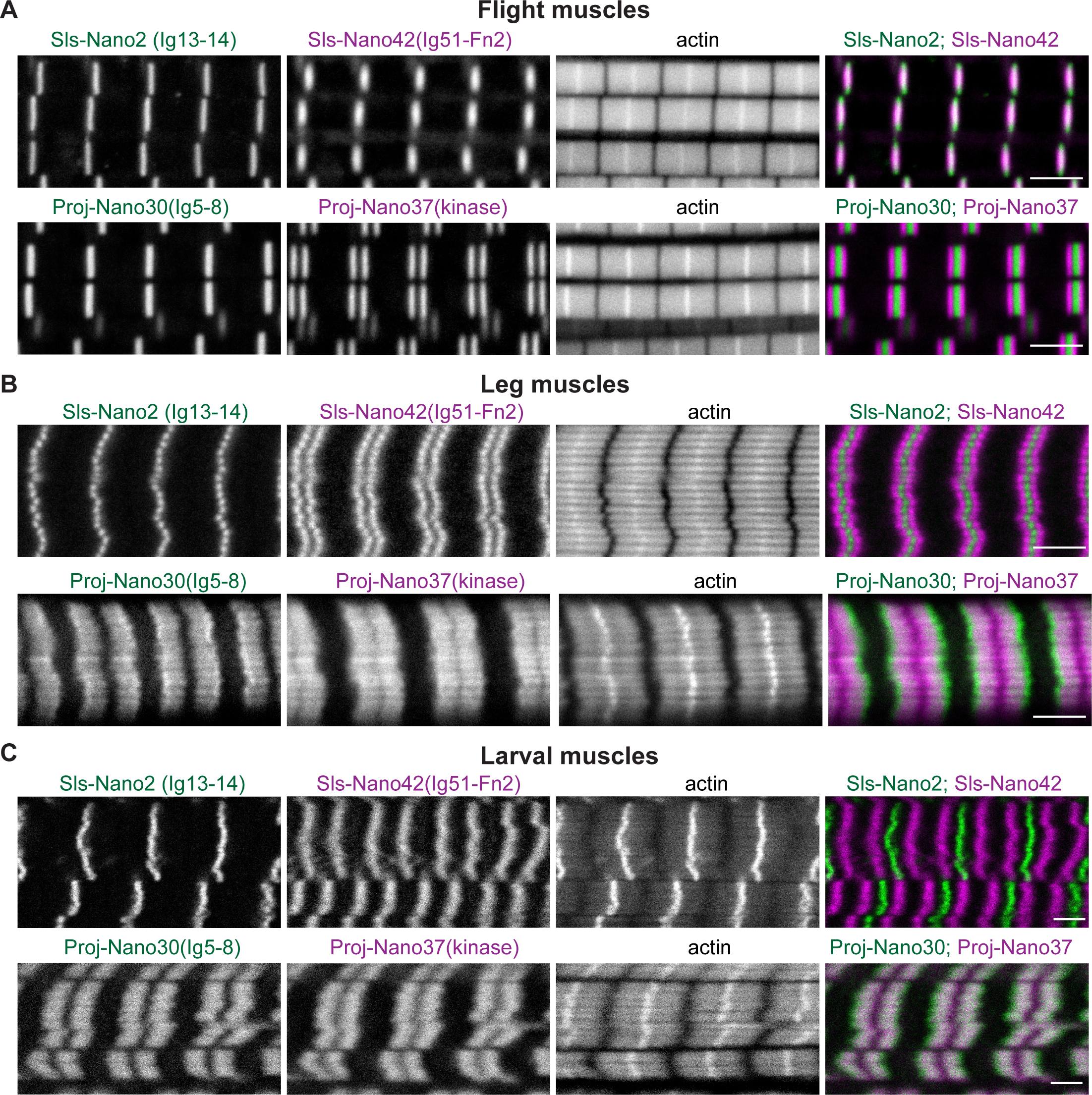
Sallimus and Projectin localisation in mature sarcomeres. (**A-C**) Mature sarcomeres from wild type flight muscles (A), leg muscles (B) or L3 larval muscles (C) stained by phalloidin (actin) together with N- and C-terminal Sls nanobodies (Sls-Nano2 in green and Sls-Nano42 in magenta, top) or N- and C-terminal Proj nanobodies (Proj-Nano30 in green and Proj-Nano37 in magenta, bottom). Scale bars are 3 µm. Note the long distance between the Sls-Nano42 bands in larval muscles and the distinct locations of Proj-Nano30 and Proj-Nano37 in leg and larval muscles.

Next, we investigated the localisation of Proj in flight muscles and found the N-terminal Ig5-8 recognised by Proj-Nano30 resulted in a single band overlapping with the Z-disc, whereas the Proj-Nano37 recognising the Proj kinase domain at its C-terminal end resulted in 2 bands right and left of the I-band, likely overlapping with the myosin filament (Figure 6A). Hence, we verified with our nanobodies that Proj is oriented in a linear fashion in flight muscles with its N-terminus closer to the Z-disc and its C-terminus facing the myosin filaments. Quantifying the precise positions of the Sls and Proj domains bound by our nanobodies in flight muscles requires super-resolution microscopy, which is reported in the accompanying manuscript (Schueder et al., 2022).

In contrast to indirect flight muscles, *Drosophila* leg or larval muscles have longer I-bands, likely caused by the expression of longer Sls splice isoforms that include large parts of the flexible PEVK spring domains making these muscles softer (Burkart et al., 2007; Spletter and Schnorrer, 2014). However, the precise positions of N- and C-terminal ends of Sls in these muscle types had not been investigated in detail prior to our studies. To address this open question, we prepared fixed adult hemi-thoraces and third instar (L3) larval filets and stained leg or larval body muscles with the N- and C-terminal Sls nanobodies. We found that Sls-Nano2 overlaps with the Z-disc in leg and larval muscles, similar to flight muscles. However, the C-terminal Sls-Nano42 recognising Sls-Ig51-Fn2 revealed 2 distinct bands with larger distances in larval muscles compared to leg muscles (Figure 6B, C). This demonstrated that *Drosophila* Sls is extended as a linear molecule bridging from the Z-disc likely to the myosin filament in sarcomeres with long I-bands.

In contrast to its defined location in flight muscles, earlier studies using Projectin antibodies suggested that Projectin largely decorates the thick filament in *Drosophila* leg muscles (Lakey et al., 1990; Saide et al., 1989; Vigoreaux et al., 1991). Consistent with these previous reports, staining of adult leg or larval body muscles with our N- and C-terminal Proj nanobodies, Proj-Nano30 and Proj-Nano37 respectively, revealed two large blocks, instead of sharp bands, located on the myosin filament in both adult leg and larval body muscles (Figure 6B, C). When carefully analysing the overlap of Proj-Nano30 and Proj-Nano37 staining, we surprisingly found that these blocks were slightly shifted in respect to each other. N-terminal Proj-Nano30 staining located closer towards the Z-disc, whereas the C-terminal Proj-Nano37 staining was closer towards the M-band (Figure 6B, C). These results suggested that Proj decorates the myosin filament of cross-striated *Drosophila* muscles in a polar fashion.

### Sallimus is stretched across long I-bands

To quantify the precise length of Sls in relaxed larval muscle sarcomeres we measured the distances between the Sls-Nano2 and Sls-Nano42 peaks. We found that Sls extends over more than 2 µm in relaxed L3 sarcomeres that are about 8.5 µm long (Figure 7A). We verified the Sls length by staining with a second Sls nanobody close to the Sls C-term, Sls-Nano39 recognising Sls-Ig49/50 (Figure 7 supplement 1A). To test if the Sls C-terminus can indeed reach the beginning of the myosin filament, we co-stained larval muscles with N- and C- terminal Sls nanobodies together with an Mhc antibody (Figure 7B). Indeed, we found that Sls-Nano42 recognising Sls-Ig51-Fn2 localises to the beginning of the myosin filaments, demonstrating that each long Sls isoform indeed stretches across the entire long I-band of larval muscles, likely to mechanically link the Z-discs to the myosin filaments.

**Figure 7.**
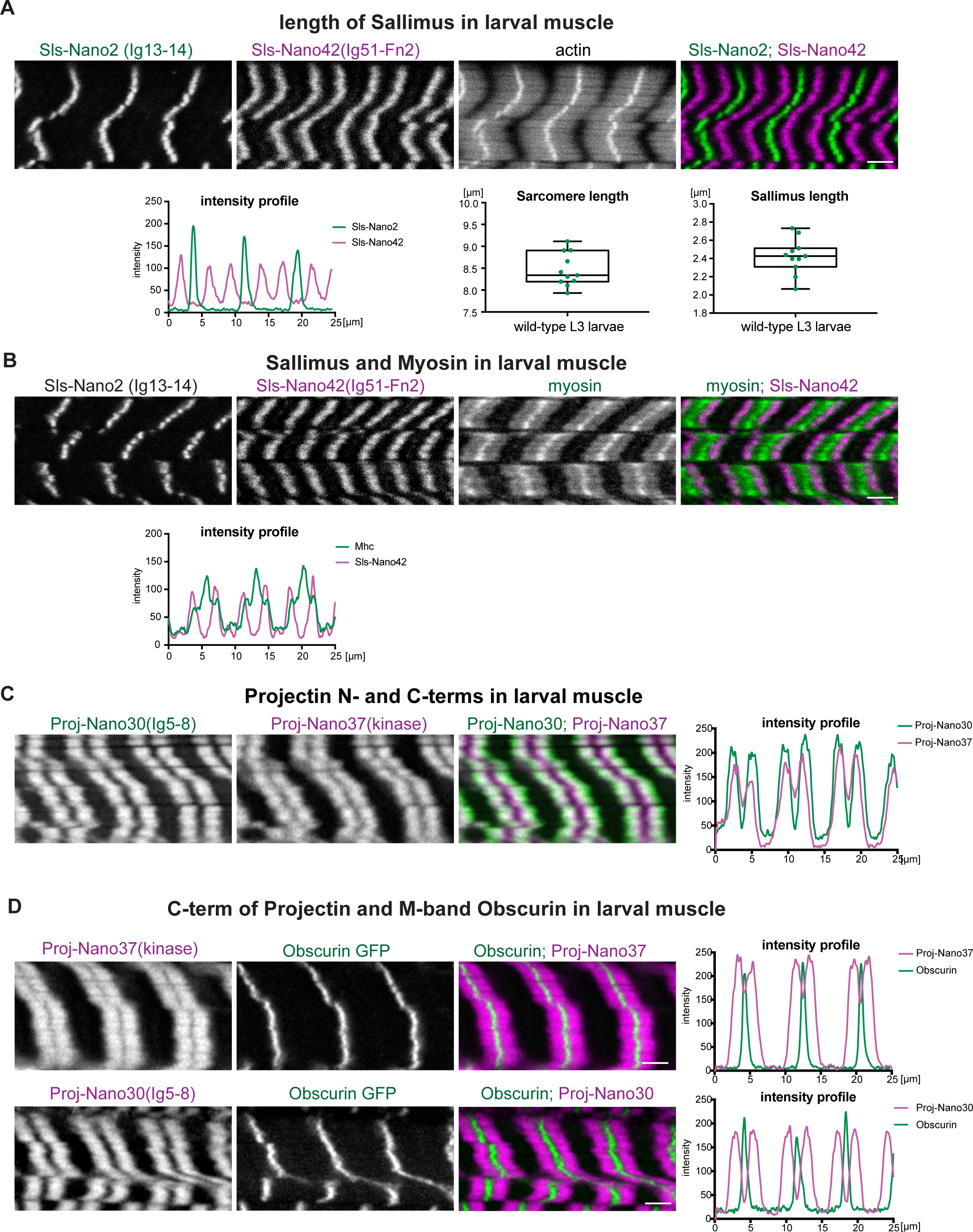
Sallimus and Projectin localisation in mature larval muscles. **(A)** L3 larval muscles stained for actin, as well as N-(Sls-Nano2, green) and C-terminal (Sls- Nano42, magenta) Sls nanobodies. Scale bar 3 µm. Plot displays longitudinal intensity profile of Sls-Nano2 and Sls-Nano42. Quantification of sarcomere length (distance between Sls- Nano2 bands) and Sls length (distance between Sls-Nano2 and Sls-Nano42). Each point represents an animal, n = 10. **(B)** L3 larval muscle stained for myosin (green), N-(Sls-Nano2) and C-terminal (Sls-Nano42, magenta) Sls nanobodies. Scale bar 3µm. Plot displays intensity profile of myosin and Sls-Nano42. Note that peaks of Nano42 map to the start of the myosin signal. **(C)** L3 larval muscle stained for Projectin with N-(Proj-Nano30, green) and C-terminal (Proj-Nano37, magenta) nanobodies and imaged with the airy-scan detector. Scale bar 3µm. Plot displays intensity profile of Proj-Nano30 and Proj-Nano37. Note that the Proj-Nano37 signal closer to the M-Band. **(D)** Obscurin GFP larvae stained for Projectin with N-(Proj- Nano30) or C-terminal (Proj-Nano37) nanobodies and imaged with an airy scan detector. Scale bar 3µm. Plots display intensity profiles. Note that Obscurin perfectly fills the Proj- Nano37 gap at the M-band.

### Projectin is oriented on the thick filament

We wanted to verify our surprising finding that N- and C-terminal Proj nanobodies result in distinct localisation patterns in sarcomeres of larval muscles. We double stained larval muscles with additional combinations of N- and C-terminal Proj nanobodies, namely Proj- Nano28 recognising Proj-Fn1/2 with Proj-Nano34 recognising the Proj kinase domain and Proj-Nano29 recognising Proj-Fn1/2 combined with Proj-Nano35 also recognising the Proj kinase domain. Again, we found that both nanobody combinations label two blocks located on the myosin filament, with Proj-Fn1/2 located closer to the Z-disc and the Proj kinase domain located closer to the M-band (Figure 7 – supplement 1B, C). Furthermore, we obtained the same result with a fourth combination of nanobodies, Proj-Nano29 recognising Proj-Fn1/2 and Proj-Nano33 binding to Proj-Ig27-Fn35 (Figure 7 – supplement 1D). This ‘shifted-blocks’ pattern is not a technical artefact as double staining with Proj-Nano30 and Proj-Nano28 or with Proj-Nano35(kinase) and Proj-Nano46(kinase) showed an almost perfect overlap (Figure 7 – supplement 1E, F). Finally, we confirmed the ‘shifted-blocks’ pattern by imaging the Proj-Nano30(Ig5-8) and Proj-Nano37(kinase) patterns with an airy-scan detector that slightly increases the spatial resolution (Figure 7C).

We hypothesized that the small central gap visible in the Proj-kinase domain nanobody patterns is caused by a Projectin-free M-band of the larval sarcomere. Co-labelling the M-band with Obscurin-GFP, which specifically localises to the M-band (Katzemich et al., 2012; Sarov et al., 2016), confirmed that the gap present in the Proj-kinase nanobody pattern is consistent with its absence from the M-band (Figure 7D). Taken together, our results demonstrate that Projectin decorates the myosin filament in a defined polar orientation, likely from the tip of the myosin filament until the beginning of the H-zone that is devoid of myosin heads.

### Live imaging of Sls using nanobodies *in vivo*

Nanobodies have the particular advantage that they are single chain proteins that can be expressed in the cytoplasm of eukaryotic cells. To our knowledge only nanobodies against GFP, mCherry or short epitope tags had thus far been expressed in *Drosophila* tissues (Caussinus et al., 2011; Harmansa and Affolter, 2018; Harmansa et al., 2015; 2017; Xu et al., 2022). Hence, we wanted to test if our nanobodies are useful tools to track a native sarcomeric protein in the mature muscle, similar to a direct GFP fusion to the sarcomeric protein. For proof of principle experiments, we chose Sls-Nano2 for two reasons: first, Sls is likely stably incorporated into mature sarcomeres and its large size should prevent fast diffusion. Thus, Sls is a suitable protein to test if a nanobody would be stably bound to a target protein in muscle. Second, we had verified the high affinity of Sls-Nano2 to the Sls-Ig13/14 target *in vitro* (Figure 3B).

To test how stably Sls-Nano2 binds to Sls *in vivo*, we expressed Sls-Nano2-Neongreen driven by *Mef2*-GAL4 during all stages of muscle development and assayed muscles of intact living L3 larvae. We found the expected striated pattern of Sls-Nano2-Neongreen (Figure 8A) resembling the Sls-Nano2 staining in fixed larval muscles. Thus, Sls-Nano2-Neongreen binds to Sls-Ig13/14 *in vivo*. To quantify the diffusion and local turn-over of Sls-Nano2-Neongreen, we established a protocol that allowed us to image intact living larvae under the spinning disc microscope for at least 30 min (see Methods). This enabled us to measure fluorescence recovery after photobleaching (FRAP) of Sls-Nano2-Neongreen in living larval muscles. We bleached one area in L3 larval muscles and measured fluorescence recovery over 29 min (Figure 8A, B, Figure 8 - Video 1). We found little to no recovery during the observation period. This demonstrates that the Sls-Nano2 is indeed stably bound the Sls-Ig13/14 target and furthermore that Sls protein does not exchange significantly over a 30 min period in mature larval muscles. Together, these data verified that nanobodies against *Drosophila* proteins can indeed be expressed *in vivo* and thus can be used to investigate the dynamics of a chosen target domain. Hence, the here generated nanobodies will be invaluable tools to quantify the dynamics of Sls and Proj during muscle development and maintenance.

**Figure 8.**
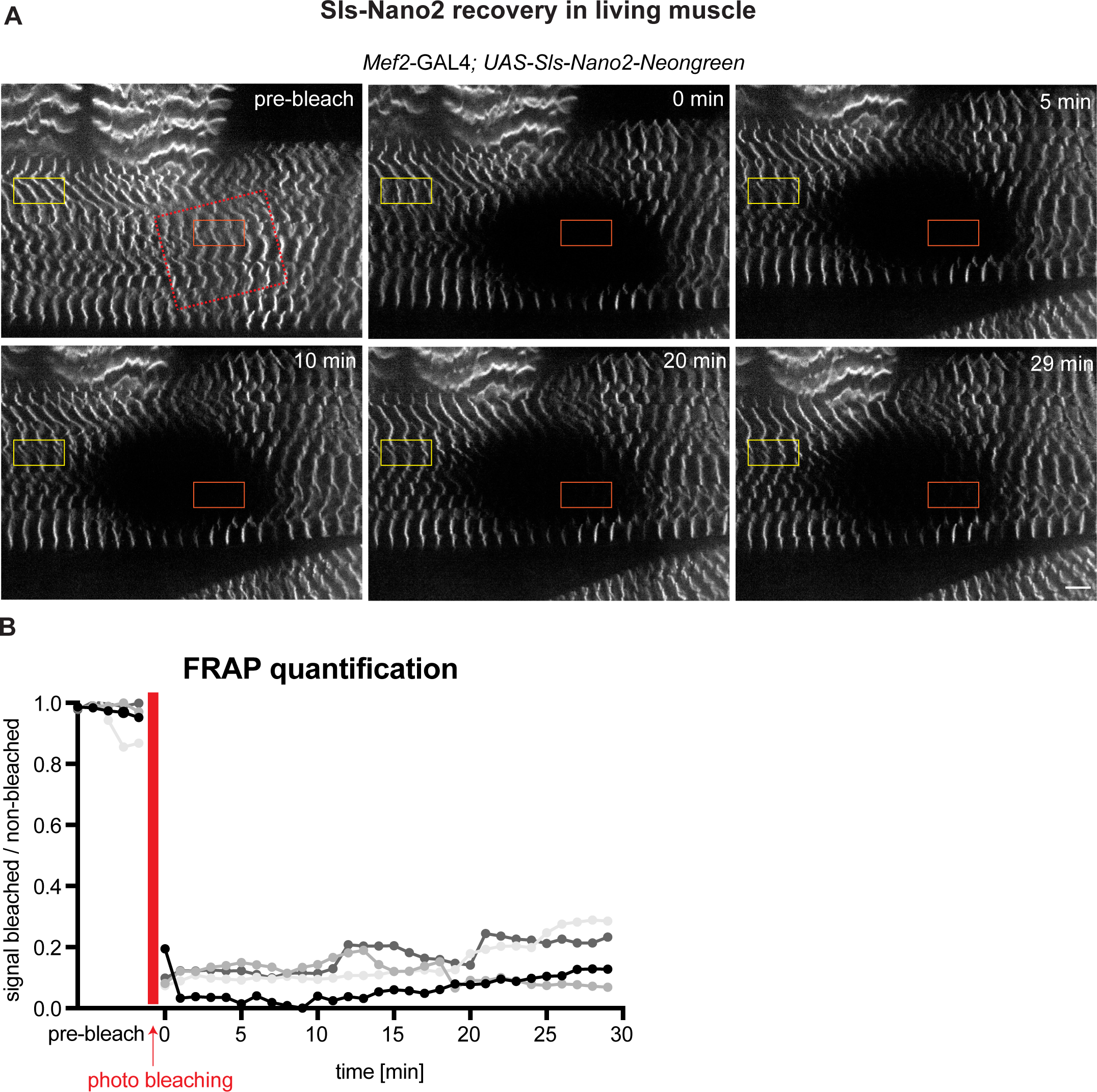
Live imaging of Sls using nanobodies *in vivo*. **(A)** Living L3 larval muscles expressing Sls-Nano2-Neongreen driven with *Mef2*-GAL4. Note the striated pattern marking the Z-disc. A region marked by the red rectangle was bleached (the larvae was slightly moving while being bleached) and fluorescence recovery was imaged. A single z-plane of a stack is shown. **(B)** Quantification of fluorescence recovery in the orange box, which was normalised by the fluorescence in the yellow box outside the bleached area. The different grey values indicate four different larvae from four different experiments. Note either absence or less than 20% recovery in the bleached area over 30 minutes.

## Discussion

### Nanobodies as tools for developmental biology

Thus far, the application of nanobodies in *Drosophila* was limited to nanobodies against fluorescent proteins or recently against short epitope tags (Caussinus et al., 2011; Harmansa and Affolter, 2018; Harmansa et al., 2015; 2017; Xu et al., 2022). These former studies had shown that nanobodies against GFP can be used to trap secreted Dpp in the *Drosophila* wing disc, and hence demonstrated the strong binding of nanobodies to their target also *in vivo* (Harmansa et al., 2015; 2017). Here we demonstrated that the high affinity of nanobodies to their targets *in vivo* is not limited to the commercially available GFP nanobody that the fly community has extensively used in the past (Caussinus et al., 2011; Harmansa and Affolter, 2018). It is likely also the case for most of the here presented Sls and Proj nanobody toolbox, as exemplified in detail for Sls-Nano2. This is significant as many GFP fusion proteins do not retain full functionality, as reported not only for Dpp-GFP but also for sarcomeric proteins such as Mhc-GFP, Sls-GFP or troponin-GFP fusion attempts (Matsuda et al., 2021; Orfanos et al., 2015; Sarov et al., 2016).

Nanobodies not only allow to monitor the dynamics of the target protein *in vivo*, as shown here for Sls, they can also be used to induce the degradation of the target protein, as shown for GFP-tagged proteins that can be degraded with the deGradFP system using a degradation box fused to the GFP nanobody (Caussinus et al., 2011; Nagarkar-Jaiswal et al., 2015). Nanobodies can also be useful as conditional blockers of their target domains, such as blocking a kinase domain as shown for EGFR in cell culture (Tabtimmai et al., 2019). Hence, the here generated nanobody toolbox can be a first step towards modulation of Sls or Proj domains activity *in vivo*.

The small size of nanobodies not only allows superior penetration into tissues as shown here for late stage *Drosophila* embryos, but also places possible labels very close to their target epitopes. This is relevant for super-resolution microscopy that can resolve the target location with a precision better than 5 nm resolution (Schnitzbauer et al., 2017) or for cryo-electron-tomography, with which the native structure of titin in the sarcomere might be resolvable in the future (Wang et al., 2022; 2021). High labelling density and proximity of the label to the target are key to identify the nature of unknown protein densities in tomograms. Hence, our toolbox should not only provide a resource to mechanistically study the function of the *Drosophila* Sls and Proj proteins in more detail in the future but may also inspire the *Drosophila* community to invest more into the generation of nanobodies in future, instead of generating antibodies by default.

### A *Drosophila* titin nanobody toolbox

We introduced here the generation and characterisation of 12 different nanobodies that were raised against 7 different target domains, 3 are present in Sls and 4 in Proj. We characterised their specificity in embryonic and larval muscles and verified that nanobodies are indeed well suited to diffuse into dense muscle tissues. They even label muscles of late stage embryos, which are impermeable to antibodies because of their chitin skeleton (Moussian, 2010).

Staining larval, leg and flight muscles with our nanobodies confirmed the existence of different Sls and Proj isoforms in the different muscle types. The stiff flight muscles do contain a short version of Sls, that does not allow to resolve the N- and C-terminal ends of Sls using confocal microscopy. This was only possible by using super-resolution microscopy with our here developed nanobodies (Schueder et al., 2022).

Our data suggest that most of the Sls isoforms present in flight muscles do contain the C-terminal Sls-Ig51-Fn2 domains. This is consistent with developmental transcriptomics results that included splice isoform annotations (Spletter et al., 2015; 2018), and the very low expression of a Sls isoform that uses an early alternative stop codon, which is rather expressed in leg muscles (Sarov et al., 2016). This is significant as the initially proposed short Sls isoform named Kettin is not supposed to contain the C-terminal Sls-Ig51-Fn2 domains and hence would not bridge across the thin I-band of flight muscles to the myosin filament (Burkart et al., 2007; Lakey et al., 1993; Szikora et al., 2020). Our new nanobodies now clarify that most Sls isoforms have at least the potential to bridge to the myosin filament in flight muscles (Schueder et al., 2022).

Similarly, our Proj nanobodies verified the defined orientation of Proj in flight muscle sarcomeres with its N-terminus facing the Z-disc and its C-terminal kinase domain oriented towards the center of myosin filament. In the accompanying manuscript these tools enabled the determination of the precise position of the Proj ends in the flight muscles using super-resolution microscopy (Schueder et al., 2022).

### A long stretched Sls isoform in larval muscle

Larval muscles are considered soft compared to stiff flight muscles. This is consistent with their large dynamic length range: larval sarcomeres have a relaxed length of about 8.5 µm and can contract up to about 4.5 µm. In contrast, flight muscle sarcomeres only contract 3.5% of their length during flight, about 120 nm (measured in *Drosophila virilis* (Chan and Dickinson, 1996)). Consistently, the I-band of relaxed larval muscles is long, about 2 µm. Hence, our finding that Sls has a length of more than 2 µm in relaxed larval muscles is only logic, considering that Sls needs to elastically bridge from the Z-disc to the myosin filament. However, this finding still comes as a surprise, since the mammalian titin is considered to be the ‘longest’ protein in the animal kingdom, about 1.5 µm long in 3 µm long relaxed human sarcomeres (Linke, 2018; Llewellyn et al., 2008; Regev et al., 2011). Mammalian titin is certainly the largest by molecular weight (up to 3800 kDa) (Brynnel et al., 2018), whereas the longest predicted *Drosophila* Sls isoform weights only 2100 kDa. Our data make it likely that *Drosophila* Sls is under strong molecular tension in larval muscles, however direct evidence by molecular tension measurements is still missing, and hence its long PEVK spring domains are likely unfolded to allow bridging of the long I-band in the relaxed state of the larval muscle (model in Figure 9). Such, Sls may store a significant amount of energy for the next round of muscle contraction, purely by unfolding its PEVK domains and not necessarily needing to unfold any of its Ig domains, which has recently been suggested for mammalian titin (Rivas-Pardo et al., 2016; 2020). Hence insect Sls might be the ‘longest’ protein naturally occurring in animals, a truly deserving member of the titin protein family.

**Figure 9.**
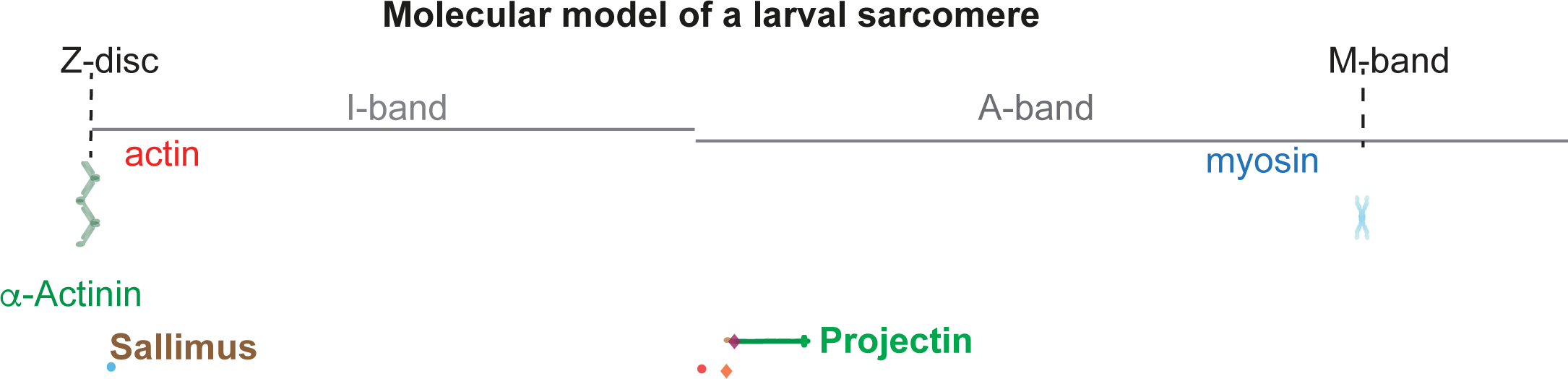
Hypothetical model of a larval sarcomere. Cartoon model of Sallimus and Projectin arrangements in a larval sarcomere. Our data reveal the polar orientation of Projectin on the myosin filament however, the precise arrangement of the individual molecules remains to be determined.

### A defined orientation of Projectin on myosin filaments

In contrast to Sls, Proj does not locate in a sharp band in larval or leg muscles, but rather as a broad block, which had been previously reported (Lakey et al., 1990; Saide et al., 1989; Vigoreaux et al., 1991). Our data revealed that the N- and C-terminal ends of Proj display slightly shifted localisations, with the C-termini located closer the M-band compared to the N-termini. This strongly suggests that each Proj protein has a defined orientation on the myosin filament (model in Figure 9). Currently, it remains unclear if neighbouring Proj molecules overlap or if they are arrayed in a linear way to decorate the thick filament, similar to how the mammalian titin protein is supposed to decorate it (Tonino et al., 2017). Super-resolution imaging of larval muscles using our nanobodies will be needed to solve this interesting question.

The fact that Proj decorates the entire thick filament of likely all *Drosophila* muscles, except indirect flight muscles, has the interesting consequence that the Proj kinase is also located along the entire thick filament. Titin kinases including the Proj kinase are proposed to be regulated by mechanical stretch: an inhibitory C-terminal tail needs to be pulled out of the kinase domain to allow kinase activity (Gautel, 2011; Gräter et al., 2005; Kobe et al., 1996; Lange et al., 2005). Thus, the larval muscle localisation of Proj would allow Proj to respond to stretch with kinase activation along the entire thick filament, and not only at the M-band as is the case for mammalian muscle. Hence, it will be interesting to resolve in the future if the kinase activity is required for sarcomere formation or function. The *Drosophila* larval muscles would be a good model to test this interesting question, and our here generated nanobodies, 4 of which target the Proj kinase domain might be a very valuable tool for these studies.

In summary, our *Drosophila* titin nanobody toolbox provides a rich resource for future studies of muscle development and maintenance in *Drosophila* and may also provide a paradigm for a strategy to investigate the molecular function and localisation of other large protein complexes during tissue morphogenesis.

## Methods

### Recombinant immunogens and Nanobody generation

We screened existing transcriptomics data (Spletter et al., 2015; 2018) and Flybase (http://flybase.org/reports/FBgn0086906; http://flybase.org/reports/FBgn0005666) to identify candidate domains of Sls and Proj that should be expressed in all muscle types. Next, we used Swissmodel (Waterhouse et al., 2018) to predict domain borders for stably folding fragments. These fragments were then codon-optimised for expression in *E.coli* and cloned into a His14- bdSUMO fusion vector (Frey and Görlich, 2014a). Expression was in *E.coli* NEB Express I^q^ at 21°C, in 2YT + 50 µg/ml kanamycin with four hours of induction with 100 µM IPTG. Bacteria were pelleted by centrifugation, resuspended in 50 mM Tris/HCl pH 7.5, 20 mM imidazole/ HCl pH 7.5, 300 mM NaCl, lysed by a freeze-thaw cycle followed by sonication. The lysate was cleared by ultracentrifugation in a T645 rotor (Thermo) at 35.000 rpm for 90 minutes. Purification by Ni(II) chelate capture and elution with 100 nM of the tag-cleaving bdSENP1 protease was as previously described (Frey and Görlich, 2014b). 100 µg of each antigen (in PBS) were used per immunisation with 200 µl Fama as an adjuvant (Gerbu #3030), following two pre-immunisations with myofibrils isolated from flight muscles of 500 adult flies.

Blood sampling, lymphocyte isolation, and construction of an M13 phage display library were done as described previously (Pleiner et al., 2015; 2018). Phage display itself was performed with 1 nM biotinylated baits immobilised to streptavidin magnetic beads. Selected clones were sequenced in a 96-well format. Coding sequences were cloned for expression into H_14_-NEDD8 or His_14_-ScSUMO vectors, with ectopic cysteines at N- and C-termini of the nanobody. The here described expression constructs will be made available at Addgene (http://www.addgene.org).

### Nanobody expression, purification, and labelling

Nanobodies were expressed in NEB Shuffle Express, which allows the structural disulphide bond to be (partially) formed. Bacteria were grown initially in 5-liter flasks containing 250 ml TB medium supplemented with 50 µg/ml kanamycin and 0.5% glucose overnight at 37°C to stationary phase (OD_600_ ∼10). The cultures were then shifted to 21°C, diluted with 500 ml fresh medium, and induced 20 minutes later with 100 µM IPTG for 4 hours.

Bacteria were pelleted, resuspended in 50 ml sonication buffer (50 mM Tris/HCl pH 7.5, 20 mM imidazole/ HCl pH 7.5, 300 mM NaCl, 5 mM GSH (reduced glutathione), 2.5 mM GSSG (oxidised glutathione)). Lysis was done by one freeze-thaw cycle followed by sonication and ultracentrifugation as described above. The lysates were then either frozen in aliquots and stored at -80°C until further use or used directly for large-scale purification. For the latter, 30 ml of lysate were bound at 4°C to 2 ml Ni(II) matrix; the matrix was extensively washed with sonication buffer, followed by protease buffer (50 mM Tris/HCl pH 7.5, 20 mM imidazole/ HCl pH 7.5, 300 mM NaCl, 5 mM GSH, 5% w/v glycerol). Elution was done with 50 nM ScUlp1 in protease buffer overnight at 4°C or for 2 hours at room temperature. Typical yields range between 10 to 50 mg nanobody per liter of culture.

For labelling, we used two different strategies. For in-solution-labelling, we reduced pre-purified nanobodies for 5 minutes with 20 mM DTT on ice. Then, free DTT was removed by gelfitration on a Nap5 Sephadex G25 column (Cytiva) equilibrated and degassed in 50mM potassium phosphate pH 6.8, 300 mM NaCl, 1 mM imidazole (using a sample volume not exceeding 400 µl). Fluorophore-maleimides were dissolved to 10 mM in dimethylformamide, used in ∼50% excess over cysteines to be labelled, pipetted into Eppendorf tubes (placed on ice) before the reduced nanobodies were added. The labelling reaction is fast and typically completed within a few minutes. Free fluorophore was then removed by gel filtration on a Nap5 column, equilibrated either in 50 mM Tris/ HCl pH7.5, 300 mM NaCl, 10% glycerol (for nanobodies with a negative net charge) or with 100 mM potassium phosphate pH 6.8, 10% glycerol (for nanobodies with a positive net charge). For storage at 4°C, 0.05% sodium azide was added. Long-term storage was at -80°C.

Quality control was done by SDS-PAGE. For most fluorophores, unlabelled, single and double labelled nanobodies are well resolved, which allows assessing the completeness of the labelling reaction (see Figure 2A). Fluorescence images were acquired from unstained/unfixed gels with a Fuji FLA-9000 system. Concentrations of nanobody, fluorophore, and density of labelling (DOL) were measured photometrically at 280 nm and at the absorption maximum of the used fluorophore. ε280 of the nanobody was deduced from its amino acid composition and used to calculate the protein concentration, also considering the cross-absorbance of the fluorophore at 280 nm. Extinction coefficients of the fluorophores at 280 nm and the absorption maximum were taken from the respective suppliers.

Alternatively, nanobodies were labelled while bound as His_14_-ScSUMO or His_14_NEDD8 fusions to a Ni(II) chelate matrix. The matrix should be resistant to reduction by DTT. We used here a homemade matrix (Frey and Görlich, 2017), however, the cOmplete his-tag purification matrix from Roche is working equally well. In brief, 30 µl Ni beads were slightly overloaded with nanobody, typically by binding 650 µl lysate to them (this usually requires titration). The beads were then washed three times in 650 µl sonication buffer; the ectopic cysteines were reduced by a 5 minutes incubation at 0°C with 20 mM DTT, 50 mM Tris/HCl pH 7.5, 300 mM NaCl, 15 mM imidazole pH 7.5. The beads were then washed twice with degassed pre-labelling buffer (50 mM potassium phosphate pH 6.8, 15 mM imidazole/ HCl pH 7.0, 300 mM NaCl). 200 µl labelling solution (100-200 µM fluorophore in 50 mM potassium phosphate pH 6.8, 1 mM imidazole/HCl pH 7.0, 300 mM NaCl) was added, the beads were shaken for 20 minutes at 0-4°C, washed twice in pre-labelling buffer, once in cleavage buffer (50 mM Tris/HCl pH 7.5, 500 mM NaCl, 20% glycerin), and finally eluted with 100 µl 50 nM ScUlp1 (in cleavage buffer) o/n at 4°C. The eluates typically contained 100 µM labelled nanobody. 50 nM was typically used for stainings.

### Biolayer interferometry (BLI)

BLI experiments were performed using High Precision Streptavidin biosensors and an Octet RED96e instrument (ForteBio/Sartorius) at 25°C with phosphate-buffered saline (PBS) pH 7.4, 0.02% (w/v) Tween-20 and 0.1% (w/v) BSA as assay buffer. Sls-Nano2, modified via an N-terminal and a C-terminal ectopic cysteine with two Biotin-PEG_3_-Maleimide molecules (Iris Biotech), was bound at 0.6 µg/ml concentration to the sensors until a wavelength shift/binding signal of 0.4 nm was reached. After one washing step in buffer, the biosensors were dipped into wells containing a concentration series of the Sls-Ig13/14 domains to measure the association rate and then incubated with assay buffer for dissociation. Data were reference-subtracted, and curves were fitted globally with a 1:1 binding model (Octet Data Analysis HT 12.0 software).

### Myofibril isolation for immunisation

We hand-dissected indirect flight muscles from 1000 adult wild-type flies from the Luminy strain (Leonte et al., 2021) in two batches of 500 each. To dissect, we cut away wings, head and abdomen and separated the thoraces into two halves along the midline using small dissection scissors (#15009-08 Fine Science Tools) and placed them into relaxing solution (100mM NaCl, 20mM NaPi pH7.2, 6mM MgCl2, 5mM ATP, 0.5% Triton X-100, complete protease inhibitor cocktail (Merck, Sigma #11697498001) with 50 % glycerol for a few minutes under the dissection scope. We then cut and scooped out the flight muscles, without taking gut or jump muscles using scissors and fine forceps (#11252-20 Dumont#5, Fine Science Tools). We collected flight muscles from 500 flies in one tube in relaxing buffer plus 50% glycerol and left them up to 24h at -20°C. Then, we spun them down the myofibrils at 200g and washed the pellet with relaxing buffer without glycerol. The purified myofibrils were then frozen in liquid nitrogen and stored at -80°C until used for alpaca immunisation.

### Fly strains and genetics

Fly stocks were maintained under standard culture conditions (Avellaneda et al., 2021). All crosses were developed at 27 °C to enhance RNAi efficiency (Schnorrer et al., 2010). Wild- type control flies were *w[1118]* or *Mef2*-GAL4 driver crossed to *w[1118]*. To knock-down *sls* or Projectin (*bt*) muscle-specific *Mef2-*GAL4 was crossed with *UAS-sls-IR* (TF47301) or *UAS-bt-IR* (TF46252) long ds-RNAi lines obtained from the VDRC stock center (Dietzl et al., 2007) and embryonic muscles were stained with nanobodies.

### Embryo fixation and staining

To investigate the larval musculature morphology at embryonic stages 16 and 17, crosses of the correct genotypes were set up in fly cages, in the presence of apple juice agar plates and a drop of yeast paste at 27°C. Flies were allowed to lay overnight and next day the embryos were collected and aged for at least another 8 h at 27°C. For fixation, embryos were dechorionated in 50% bleach for 2-3 min and then fixed for 20 min with a 1:1 mixture of 4% paraformaldehyde (PFA in fresh PBS) and heptane in glass tubes on shaker at room temperature. To free the embryos from the vitelline membrane, the fixative (lower phase) was removed with a glass pipette and one volume of methanol was added and the tube was shaken vigorously. Dechorionated embryos sunk to the bottom and were washed 3x with MeOH. Embryos were stored at -20°C in MeOH.

For antibody and nanobody stainings, embryos were rehydrated in PBST (PBS with 0.3% Triton-X-100), blocked for more than 30 min with 4% normal goat serum and stained with fluorescently labelled nanobodies alone, or together with antibodies, overnight in PBST (PBS with 0.3% Triton-X-100). Antibodies were visualised with standard secondary antibodies (Molecular Probes, 1/500 in PBST) embryos were mounted in SlowFadeTM Gold Antifade (Thermofisher) and imaged with a Zeiss LSM880 confocal microscope using a 40x objective.

### Flight and leg muscle staining

Flight and leg muscles were stained as published in detail before (Weitkunat and Schnorrer, 2014). Briefly, wings, head and abdomen were clipped from adult flies with fine scissors and thoraces were fixed in 4% PFA in PBS-T (PBS with 0.3% Triton X-100) for 20 min at room temperature. After washing once with PBST, the thoraxes were placed on a slide with double-sticky tape with the head position facing the sticky tape and cut sagittally with a microtome blade (Pfm Medical Feather C35). Hemi-thoraces were stained with fluorescent nanobodies and rhodamine-phalloidin (1:1000 Molecular Probes) for 2 hrs at room temperature. Hemi- thoraces were washed twice with PBS-T, mounted in SlowFadeTM Gold Antifade (Thermofisher) using two cover slips as spacers and flight or leg muscles were imaged with a Zeiss LSM880 confocal microscope using 63x objective.

### Dissection and staining of larval muscles

To perform antibody or nanobody stainings of larval muscles, third instar (L3) larvae were collected with a brush and placed at 4°C. For dissection, larvae were covered with HL3 buffer and pinned individually by pushing one insect pin through the head and one through the abdomen to immobilize them in dissection dishes placed on ice (Stewart et al., 1994). Pinned larvae were dissected with sharp scissors from the dorsal side in HL3 buffer and interior organs (gut and fat body) were removed with forceps. The remaining larval filets were fixed in 4% PFA in PBS-T (PBS with 0.3% Triton X-100) for 30 min, and then blocked in 4% normal goat serum for 30 min at room temperature (RT) on a shaker. Nanobodies and antibodies were incubated in PBS-T overnight at 4°C. Larval filets were then washed 3 times 10 min in PBST at RT and stained with secondary antibodies and phalloidin (labelled with rhodamine 1:1000, Molecular Probes) in PBST for 2 h, at RT in the dark. After washing 3 times with PBST for 5 min, larval filets were mounted in SlowFadeTM Gold Antifade (Thermofisher) and imaged with a Zeiss LSM880 confocal microscope using 20x, 40x objectives.

To quantify larval sarcomere and Sls length, the images were processed with a Gaussian blur (sigma: 1.00) and a line perpendicular to the Z-disc was drawn to retrieve an intensity profile. The position of the peak of intensity was determined by using BAR plugin in Fiji (Schindelin et al., 2012). Sarcomere length was calculated by the distance between two peaks of Sls-Nano2 staining and Sls length by the distance between a peak of Sls-Nano2 and Sls-Nano42.

### Generation of UAS-Nano2-mNeonGreen transgenic flies

To clone *UAS-Nano2-NeonGreen*, we first linearised pUAST-attB with EcoRI and inserted mNeonGreen by Gibson Assembly (Gibson Assembly®) after amplification of mNeonGreen with 5’-ACTCTGAATAGGGAATTGGGAATTC-3’ and 5’-CGGCCGCAGATCTGTTAAC-3’ primers. In a second step, we linearised pUAST-attB- mNeonGreen with EcoRI and inserted the Sls-Nano2 sequence by Gibson Assembly (Gibson Assembly®) after amplification with 5’- ACTCTGAATAGGGAATTGGG-3’ and 5’- CCTTGCTCACCATGGAAC-3’ primers. Finally, we inserted the pUAST-attB-Sls-Nano2- mNeonGreen plasmid at attP site VK00033 by standard injection and selection methods (Sarov et al., 2016).

### Live imaging of larval muscles

To quantify Sls-Nano2 localisation *in vivo*, we crossed *UAS-Sls-Nano2-mNeonGreen* flies with *Mef2-*GAL4 and collected L3 larvae. To reduce movement of the living larvae, larvae were anaesthetised for 5 min with diethylether (Aldrich) (Kakanj et al., 2020) and then mounted in 10S halocarbon oil. Larvae were imaged with an Olympus spinning disc confocal microscope with a 60x objective. Photo-bleaching was performed with a 488 nm laser (Rapp- opto) and recovery was quantified during 30 min. Regions of interests (20 x 10 µm) inside the bleached area, in the non-bleached area or outside the muscle as background were selected and their intensities were measured at each time point. To calculate the ratio of FRAP the intensity of the bleached area background subtracted was divided by the intensity in the non-bleached area background subtracted.

### List of used antibodies and nanobodies

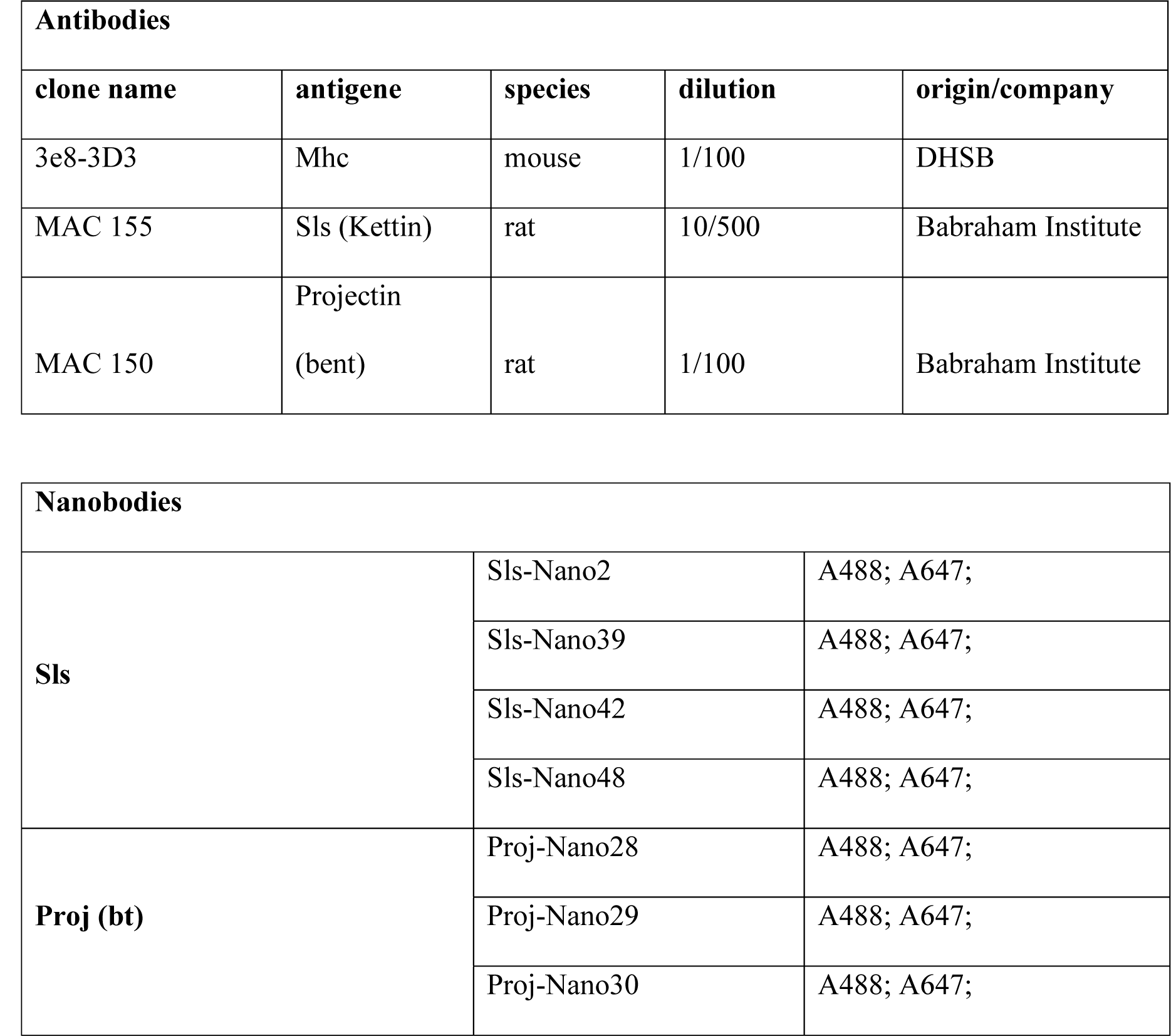
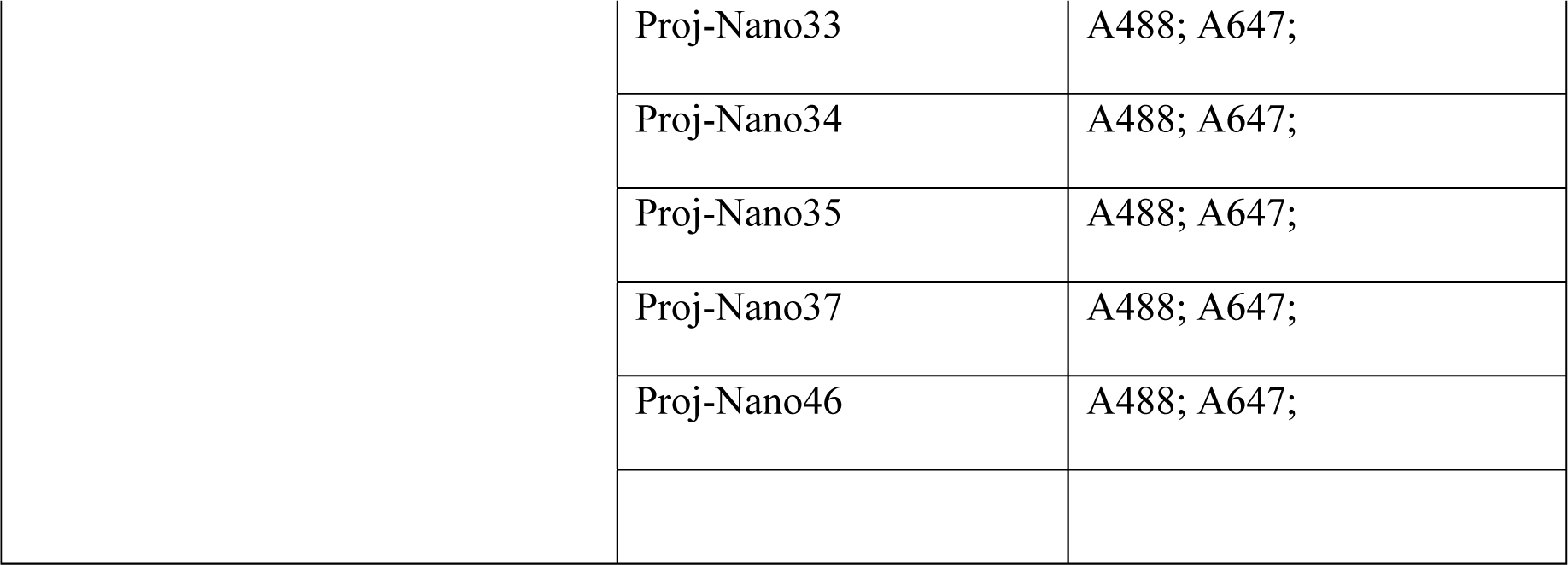

| Antibodies |  |  |  |  |
| --- | --- | --- | --- | --- |
| clone name | antigene | species | dilution | origin/company |
| 3e8-3D3 | Mhc | mouse | 1/100 | DHSB |
| MAC 155 | Sls (Kettin) | rat | 10/500 | Babraham Institute |
| MAC 150 | Projectin<br>(bent) | rat | 1/100 | Babraham Institute |

| Nanobodies |  |  |
| --- | --- | --- |
| Sls | Sls-Nano2 | A488; A647; |
|  | Sls-Nano39 | A488; A647; |
|  | Sls-Nano42 | A488; A647; |
|  | Sls-Nano48 | A488; A647; |
| Proj (bt) | Proj-Nano28 | A488; A647; |
|  | Proj-Nano29 | A488; A647; |
|  | Proj-Nano30 | A488; A647; |
|  | Proj-Nano33 | A488; A647; |
|  | Proj-Nano34 | A488; A647; |
|  | Proj-Nano35 | A488; A647; |
|  | Proj-Nano37 | A488; A647; |
|  | Proj-Nano46 | A488; A647; |

## Acknowledgements

We thank Stefan Raunser and Mathias Gautel and all their group members, as well as the Schnorrer and Görlich groups for their stimulating discussions within the StuDySARCOMERE ERC synergy grant. We thank Metin Aksu for help with the Octet measurements. We are grateful to the IBDM imaging facility for help with image acquisition and maintenance of the microscopes.

## Funding

This work was supported by the Centre National de la Recherche Scientifique (CNRS, F.S.), the Max Planck Society (D. G.), Aix-Marseille University (P.M.), the European Research Council under the European Union’s Horizon 2020 Programme (ERC-2019-SyG 856118 to D.G. & F.S.), the excellence initiative Aix-Marseille University A*MIDEX (ANR-11-IDEX- 0001-02, F.S.), the French National Research Agency with ANR-ACHN MUSCLE-FORCES (F.S.), the Human Frontiers Science Program (HFSP, RGP0052/2018, F.S.), the Bettencourt Foundation (F.S.), the France-BioImaging national research infrastructure (ANR-10-INBS- 04-01) and the Investissements d’Avenir, French Government program managed by the French National Research Agency (ANR-16-CONV-0001) and from Excellence Initiative of Aix-Marseille University - A*MIDEX (Turing Center for Living Systems) and LabEx-INFORM (F.S. & V.L.). The funders had no role in study design, data collection and analysis, decision to publish, or preparation of the manuscript.

## Competing interests

The authors declare no competing interests.

## Supplementary figures legends

**Figure 4 – figure supplement 1.**
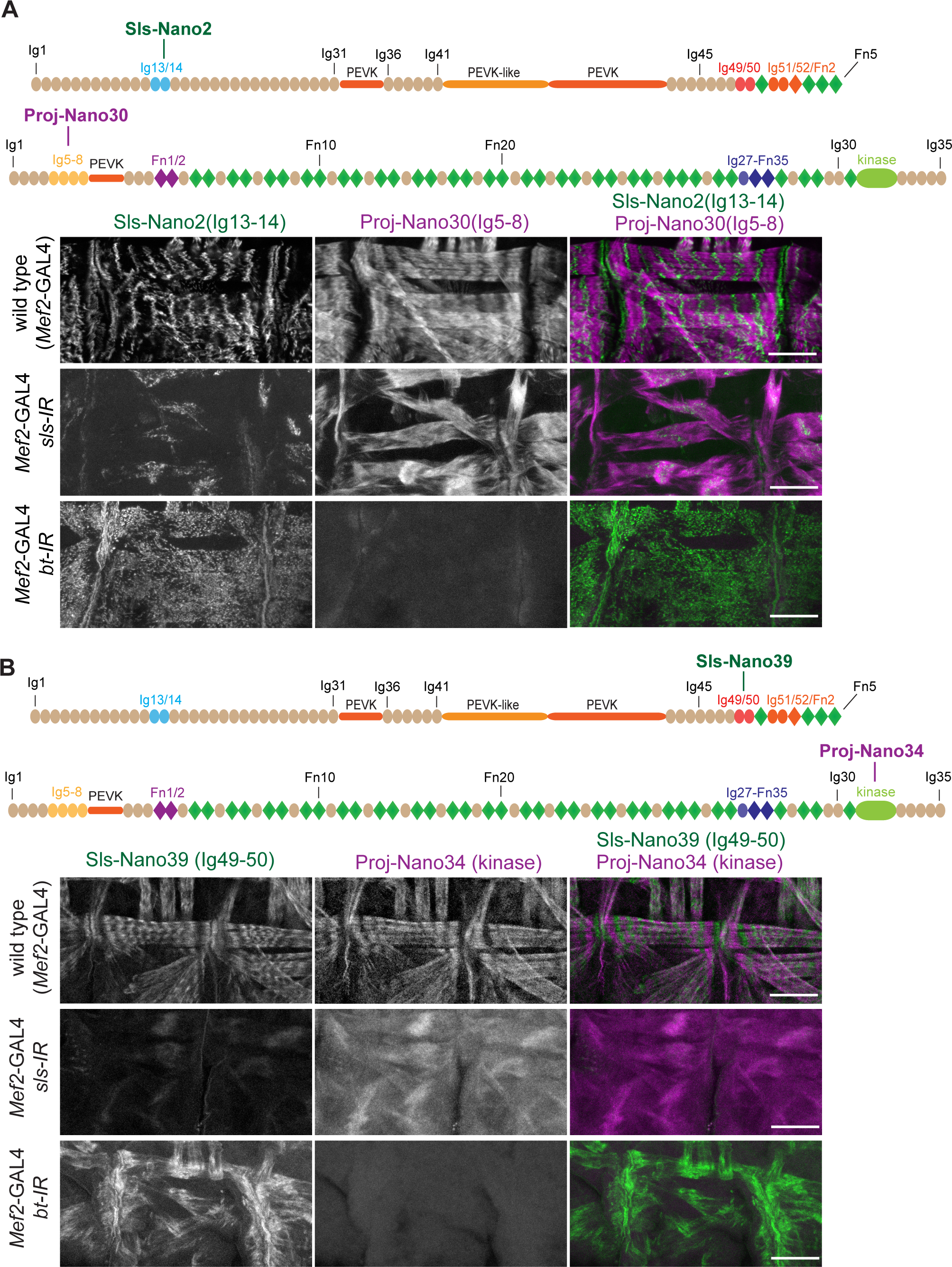
(**A, B**) top: schematic representation of Sallimus or Projectin domains with nanobodies used for stainings. Bottom: stage 17 embryos of wild type (*Mef2*-GAL4) and muscle specific *sls* or *bt* knock-down (*Mef2*-GAL4, UAS-*sls-IR* and *Mef2*-GAL4, UAS-*bt-IR*, respectively) stained with Sls (green) and Proj (magenta) nanobodies Sls-Nano2 and Proj-Nano30 (A) or Sls- Nano39 and Proj-Nano34 (B). Note the striated pattern of Sls and Proj in wild type, which is lost upon knock-down of one component. Scale bars are 20 µm.

**Figure 4 – figure supplement 2.**
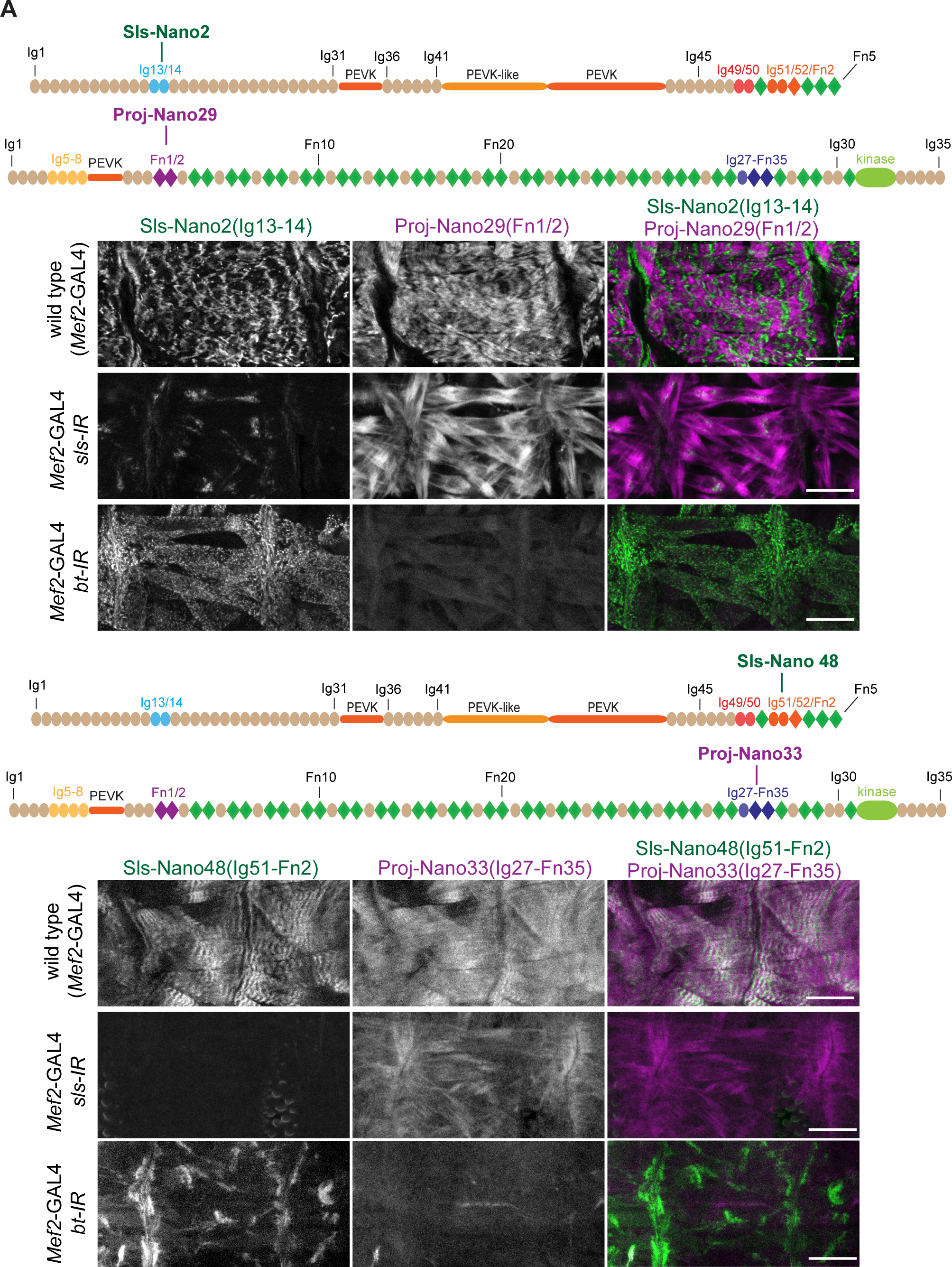
(**A, B**) top: schematic representation of Sallimus or Projectin domains with nanobodies used for stainings. Bottom: stage 17 embryos of wild type (*Mef2*-GAL4) and muscle specific *sls* or *bt* knock-down (*Mef2*-GAL4, UAS-*sls-IR* and *Mef2*-GAL4, UAS-*bt-IR*, respectively) stained with Sls (green) and Proj (magenta) nanobodies Sls-Nano2 and Proj-Nano29 (A) or Sls- Nano48 and Proj-Nano33 (B). Note the striated pattern of Sls and Proj in wild type, which is lost upon knock-down of one component. Scale bars are 20 µm.

**Figure 7 – figure supplement 1.**
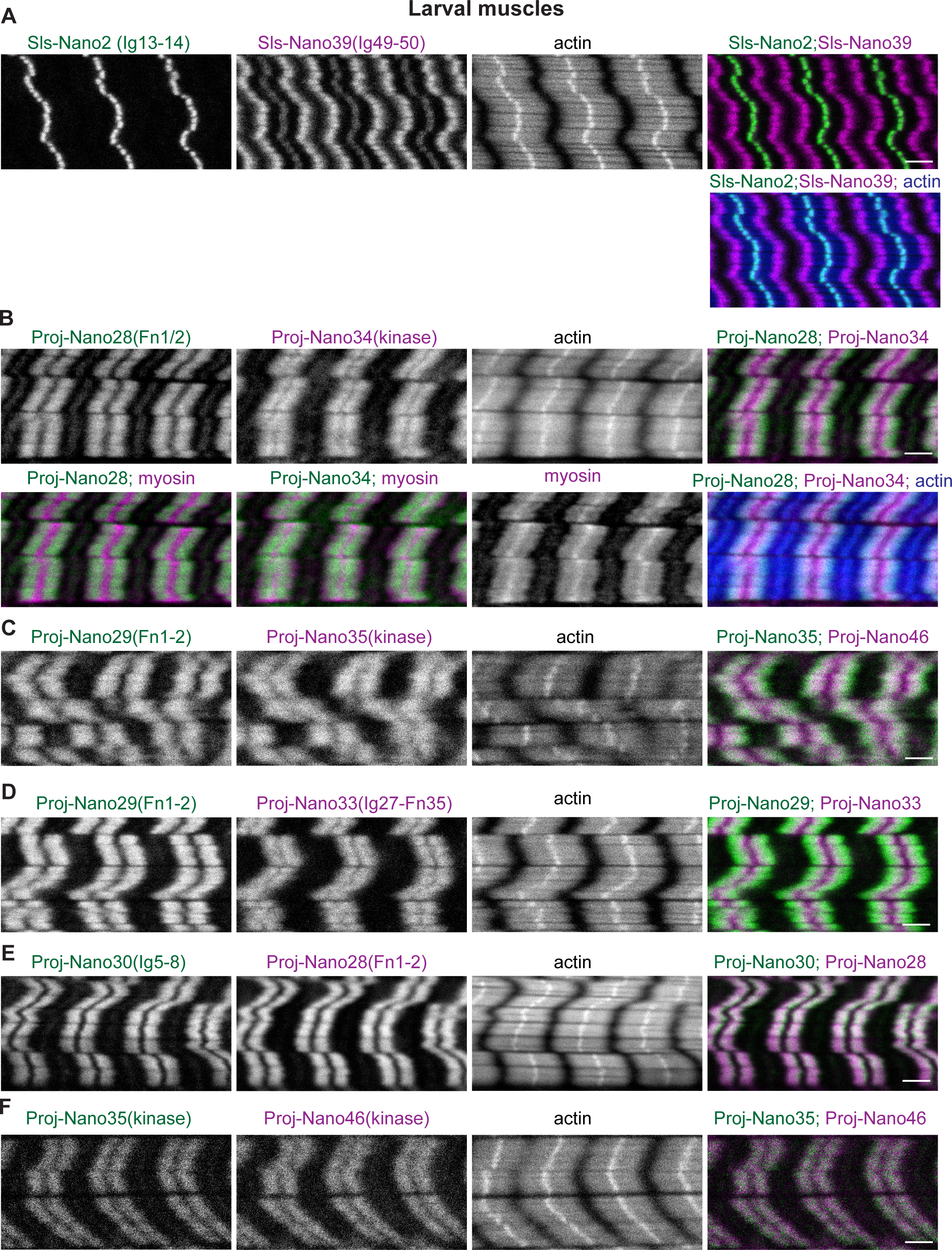
**(A)** Wild type L3 larval muscle stained for actin and Sallimus with N-(Sls-Nano2, green) and C-terminal (Sls-Nano39, magenta) Sls nanobodies. Note the large distance of the Sls-Nano39 bands (a central SlsNano39 band at the Z-disc is sometimes seen that may indicate mechanical detachment of Sls during the muscle preparation). **(B)** Wild type L3 larval muscle stained for actin, myosin and Projectin with N-(Proj-Nano28, green) and C-terminal (Proj- Nano34, magenta) nanobodies. Note the shift of N- versus C-terminal nanobodies, both label the myosin filament. **C)** Wild type L3 larval muscle stained for actin and Projectin with N- (Proj-Nano29, green) and C-terminal (Proj-Nano35, magenta) nanobodies. **(D)** Wild type L3 larval muscle stained for actin and Projectin with N-(Proj-Nano29, green) and C-terminal (Proj-Nano33, magenta) nanobodies. **(E, F)** Wild type L3 larval muscle stained for actin and Projectin with two N-terminal (Proj-Nano30, green and Proj-nano28, magenta, in E) or two C-terminal (Proj-Nano35, green and Proj-Nano46, magenta in F) nanobodies. Note that the two N-terminal or C-terminal nanobody patterns overlap. Scale bars are 3 µm.

Figure 8 – video 1. Live imaging of Sls with a nanobody using FRAP Living L3 larva expressing Sls-Nano2-Neongreen was imaged with a spinning disc confocal. One single plane is shown and assembled into the movie. The white square after the pre-bleach frames indicates the bleached area. One z-stack was recorded each minute after bleaching for 30 min, from which one frame is shown. Note that the larva is moving slightly during the experiment. There is little recovery of Sls-Nano2-Neongreen in the bleached area over a 30 min period.

## Notes

### Competing Interest Statement

The authors have declared no competing interest.

